# Identifying Novel Targets by using Drug-binding Site Signature: A Case Study of Kinase Inhibitors

**DOI:** 10.1101/860510

**Authors:** Hammad Naveed, Corinna Reglin, Thomas Schubert, Xin Gao, Stefan T. Arold, Michael L. Maitland

## Abstract

Current FDA-approved kinase inhibitors cause diverse adverse effects, some of which are due to the mechanism-independent effects of these drugs. Identifying these mechanism-independent interactions could improve drug safety and support drug repurposing. We have developed “iDTPnd”, a computational approach for large-scale discovery of novel targets for known drugs. For a given drug, we construct a positive and a negative structural signature that captures the weakly conserved structural features of drug binding sites. To facilitate assessment of unintended targets iDTPnd also provides a docking-based interaction score and its statistical significance. We were able to confirm the interaction of sorafenib, imatinib, dasatinib, sunitinib, and pazopanib with their known targets at a sensitivity and specificity of 52% and 55% respectively. We have validated 10 predicted novel targets, using *in vitro* experiments. Our results suggest that proteins other than kinases, such as nuclear receptors, cytochrome P450 or MHC Class I molecules can also be physiologically relevant targets of kinase inhibitors. Our method is general and broadly applicable for the identification of protein-small molecule interactions, when sufficient drug-target 3D data are available.

## Introduction

Proteins that contain kinase domains are involved in numerous cellular processes including signaling, proliferation, apoptosis, and survival [1,2]. The human kinome consists of more than 500 members [3]. These kinases have diverse sequences but a high degree of 3D structure similarity, particularly in the ATP binding pocket [4]. Kinases are the primary drug targets for the treatment of many cancers [5–7]. There are more than 30 FDA-approved small molecule kinase inhibitors that bind kinase domains reversibly or irreversibly. The kinase inhibitors that bind reversibly can be categorized into four major types based on the binding pocket conformation and the aspartate-phenylalanine-glycine (DFG) motif of the kinase activation loop, that controls access to the binding pocket [8,9]. Most of these inhibitors fall into the type I or type II categories. Type I kinase inhibitors bind in an ATP-competitive manner to the active forms of kinase domains with the aspartate amino acid facing into the active site. Type II kinase inhibitors on the other hand bind the inactive forms of kinase domains with the aspartate residue facing outside the active site [8,9]. Given that the ATP binding site has necessarily conserved features across most kinase domains, several kinase inhibitors interact with the human kinome broadly and are not very selective (on average, 135 or 26% of all human kinases interact with one or more kinase inhibitors included in this study) [10,11]. This broad reactivity affects the inhibitor’s efficacy and toxicity [12–14]. Therefore, predicting kinase inhibitor (off-) targets is central for the rapid and cost-efficient development of inhibitors as it allows to better understand a drug’s adverse effects and to explore drug repositioning opportunities [15].

Recent studies estimate that many unintended targets of approved drugs are yet to be discovered [16,17] and this mechanism-independent binding leads to toxicity [14]. When unexpected adverse effects of new drugs, especially kinase inhibitors are observed, it can be difficult to determine whether these effects are due to the compound binding additional unexpected targets or to a previously unknown relationship between a drug’s intended target and the function of a complete human organ system. Therefore, predicting mechanism-independent binding sites could enhance early evaluation of a compound’s specificity and hence the likelihood for specific clinical consequences.

Computational methods have increasingly been used for hit identification and lead optimization [18]. These methods fall into four categories: methods that use i) binding site structure, ii) gene expression, iii) ligand structure, and iv) a combination of the above. Structure-based methods employ binding site similarity and/or molecular docking [19–22]; expression-based methods use change in expression levels of the proteins that results from drug activity [23–27]; ligand-based methods utilize the structural and chemical properties of a drug [28–30]; and hybrid methods combine two or more types of data [31–35]. In addition, novel targets for drugs have also been identified by comparing adverse effects [36] and by using genome-wide association studies [37].

In this paper, we propose iDTPnd (integrated <u>D</u>rug <u>T</u>arget <u>P</u>redictor with <u>n</u>egative <u>d</u>ataset), a computational method for large-scale discovery of new drug targets that markedly improves our previous methodology [38] by incorporating a negative structural signature (conserved structural signatures in the kinases that are known not to interact with the respective drug). We now also provide a docking-based interaction score along with its statistical significance. In a blind test of 5 FDA-approved kinase inhibitors, we predicted the known targets with 52% sensitivity and 55% specificity. This is a significant improvement compared to a baseline model based on sequence similarity and to a recently published study [17], which reports a precision of 30% and a recall of 27% with an estimated false positive rate of 70%. In addition, our methodology is generic and can be used broadly for all types of small molecule drugs for which sufficient (~30) 3D structures of known targets are available. We have also validated 10 predicted interactions through *in vitro* experiments. It is important to note that our predictions are not limited to kinases.

## Materials and Methods

### Dataset

We extracted the positive and negative datasets from the kinome scan assay of Davis et al., 2011 [11]. Kinase inhibitors for which there was one co-crystallized structure with its target and at least 30 known targets with experimentally determined structures (apo or bound to other entities) available were selected for this study (Table 1, Supplementary Table 1). Redundancy reduction was carried out in the following manner: for the positive dataset, 70% sequence identity cut-off was used for all structures that were not bound to the respective drug and all co-crystallized structures were included (usually 1-4) in the positive dataset. For the negative dataset, as we had a relatively larger dataset we used a stricter cut-off of 60% sequence identity. We did not do redundancy reduction between the positive and the negative dataset. As structural databases are growing exponentially, the number of drugs to which the method can be applied is expected to increase significantly. The total structures deposited in PDB at the end of 2011 were 77,452 as compared to 150,593 structures on April 4, 2019 (https://www.rcsb.org/). This means that the number of structures has almost doubled in 7 years. Similarly, in case of membrane protein structures which are the targets of more than 60% marketed drugs, we had 328 structures in 2011. This number has now increased to 876 structures (https://blanco.biomol.uci.edu/mpstruc/). Therefore, we expect our method to be applicable for broader set of studies going forward.

**Table 1.** Self Validation and Comparison with Baseline. For 5 of the kinase inhibitors in our dataset, we can predict the known targets with an average of 52% sensitivity and 55% specificity. For Sorafenib we achieved the highest sensitivity and specificity of 68% and 59% respectively. Prediction using the positive signature alone is not reliable as demonstrated by the large standard deviation (given in brackets). iDTPnd performs significantly better than the baseline sequence based model in terms of sensitivity and specificity.

| Drug | iDTP |  | iDTPnd |  | Baseline_Seq_Sim |  |
| --- | --- | --- | --- | --- | --- | --- |
|  | Positive Signature |  | Positive + Negative Signature |  |  |  |
|  | Sensitivity (TP/P) | Specificity (TN/N) | Sensitivity (TP/P) | Specificity (TN/N) | Sensitivity (TP/P) | Specificity (TN/N) |
| sorafenib | 81% | 33% | 68% | 59% | 42% | 37% |
| imatinib | 3% | 94% | 55% | 54% | 27% | 26% |
| dasatinib | 2% | 99% | 42% | 57% | 32% | 29% |
| sunitinib | 63% | 67% | 46% | 50% | 48% | 38% |
| pazopanib | 4% | 95% | 50% | 55% | 32% | 37% |
| **Average (±**<br>**Sd)** | **31**<br>**($\pm 34$)%** | **78**<br>**($\pm 25$)%** | **52 ($\pm 9$)%** | **55 ($\pm 3$)%** | **36 ($\pm 8$)%** | **33 ($\pm 5$)%** |

**Table 2.**
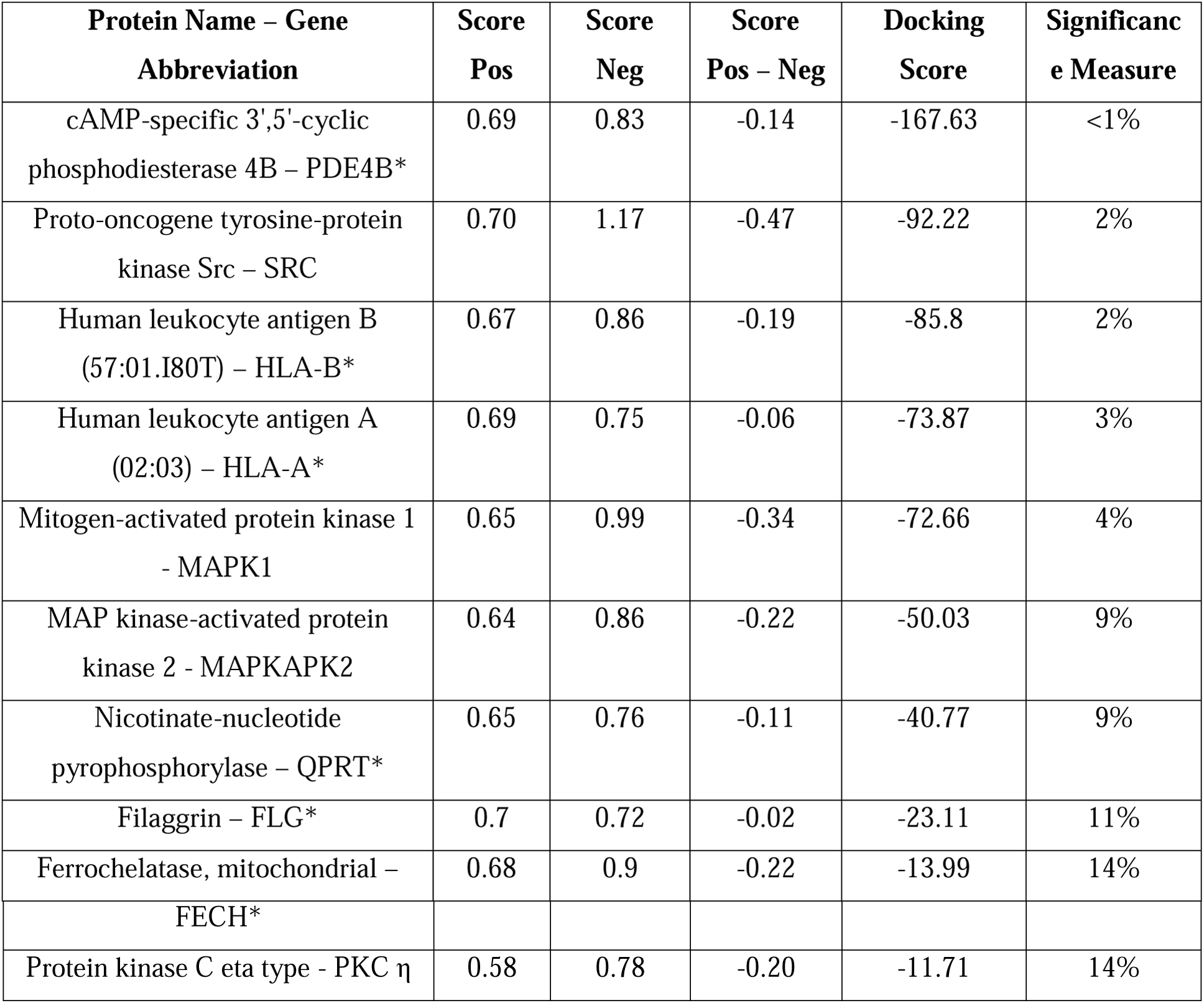
Predicted targets of Sorafenib. Top 10 predicted targets of sorafenib, their PDB id, Score_positive_, Score_negative_, Score_positive_ – Score_negative_, docking score and the significance measure (random chance to obtain a better docking score). The predicted targets had less than 60% sequence similarity with the known targets of sorafenib. For HLA-A and HLA-B, the specific alleles are given in brackets.* denotes those proteins that do not contain a kinase domain.

| **Protein Name – Gene**<br>**Abbreviation** | **Score**<br>**Pos** | **Score**<br>**Neg** | **Score**<br>**Pos – Neg** | **Docking**<br>**Score** | **Significance Measure** |
| --- | --- | --- | --- | --- | --- |
| cAMP-specific 3',5'-cyclic phosphodiesterase 4B – PDE4B* | 0.69 | 0.83 | -0.14 | -167.63 | <1% |
| Proto-oncogene tyrosine-protein kinase Src – SRC | 0.70 | 1.17 | -0.47 | -92.22 | 2% |
| Human leukocyte antigen B (57:01.I80T) – HLA-B* | 0.67 | 0.86 | -0.19 | -85.8 | 2% |
| Human leukocyte antigen A (02:03) – HLA-A* | 0.69 | 0.75 | -0.06 | -73.87 | 3% |
| Mitogen-activated protein kinase 1 - MAPK1 | 0.65 | 0.99 | -0.34 | -72.66 | 4% |
| MAP kinase-activated protein kinase 2 - MAPKAPK2 | 0.64 | 0.86 | -0.22 | -50.03 | 9% |
| Nicotinate-nucleotide pyrophosphorylase – QPRT* | 0.65 | 0.76 | -0.11 | -40.77 | 9% |
| Filaggrin – FLG* | 0.7 | 0.72 | -0.02 | -23.11 | 11% |
| Ferrochelatase, mitochondrial – | 0.68 | 0.9 | -0.22 | -13.99 | 14% |
| FECH* |  |  |  |  |  |
| Protein kinase C eta type - PKC $\eta$ | 0.58 | 0.78 | -0.20 | -11.71 | 14% |

### Sequence Similarity Baseline Model

The sequence similarity baseline model used nearest neighbor algorithm to allocate a protein to the interacting or non-interacting cluster. Using leave-one-out cross validation, global pairwise sequence similarity (not identity) was calculated between the left out protein and all other proteins. The left out protein was assigned to the cluster that contained the protein with the maximum pairwise global sequence similarity to the left out protein. If none of the protein pairs had a global sequence similarity > 0.6 then a label was not assigned to the left out protein.

### Constructing the structural signature

The flowchart of our method is shown in Supplementary Fig 1. Briefly, CASTp webserver was used to extract the pocket that the drug binds to [39], referred to as the ‘bound pocket’ from here on. Sequence order-independent alignment was used to find the pocket similar to the bound pocket [40,41] using the distance function described below. We extracted the conserved (positive and negative) structural signatures by applying pairwise sequence order-independent structure alignment followed by hierarchical clustering.

> *Score = Structural Score +* α * *Sequence Score*
>
> *Structural Score = RMSD* * *N*^(−*1/3)*^
>
> *Sequence Score = 1- (Sequence Similarity / Best Sequence Similarity)*
>
> *Sequence Similarity = ∑*_*i*_ *(AtomFreq*_*i*_ *+ ResFreq*_*i*_)
>
> *Best Sequence Similarity = ∑*_*i*_ *(MaxAtomFreq*_*i*_ *+ MaxResFreq*_*i*_)

**Figure 1.**
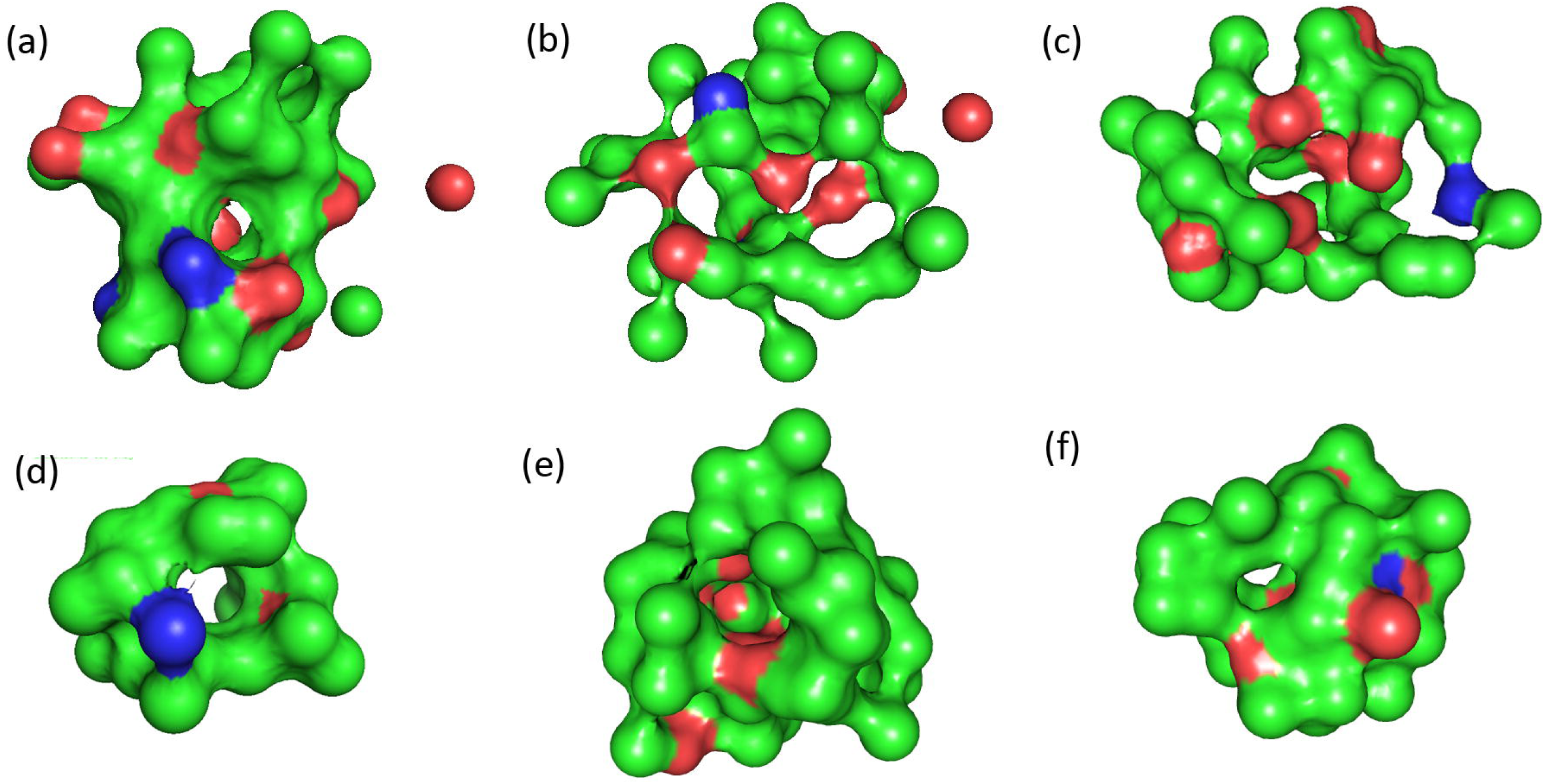
Structural Signatures. 3D structural signatures (positive, negative) of Sorafenib (a, d), Imatinib (b, e), and Dasatinib (c, f), where the color of each position represents the atom with the highest frequency. Color codes are Carbon: green, Oxygen: red, Nitrogen: blue.

Our method is not sensitive to the exact value of α as long as it is close to 1. The α can be adjusted according to empirical insight from the data. For this study we use α = 1.2. RMSD is the root mean square distance, N is the number of positions aligned, AtomFreq_i_/ResFreq_i_ represents the frequency of atom/residue aligned at position i, MaxAtomFreq_i_/MaxResFreq_i_ represents the maximum frequency of any atom/residue aligned at position i, and the summation is over all aligned positions. Every position in the signature is present in at least 50% of the structures. To achieve a minimalistic structural signature, preservation ratio cutoff is increased if the number of atoms in the signature is more than 100. While combining the positive and the negative structural signatures, predicted targets are those that have better positive score than the negative one (Score_positive_ – Score_negative_ < 0).

### MicroScale Thermophoresis

The predicted protein targets HLA-A (Acris Antibodies GmbH: item # TP300661), HLA-B (Acris Antibodies GmbH: item # TP310631), MAPKAPK2 (Merck Millipore: item # 14-337), PDE4B (Hölzel Diagnostika Handels GmbH: item # 11527-H20B-20), PKC η (abcam: item # ab60849), ERα (Biozol: item # USC-RPB050HU01-50), CDK2 (antikoerper: item # ABIN2003156), and tyrosine-protein kinase ITK/TSK (ITK) (Biomol: item # BPS-40445) were labeled by NHS chemistry with the help of an NT647-labeling kit by NanoTemper Technologies. In an initial step the Tris containing storage buffers were exchanged by the MST labeling buffer as indicated by the manufacturers in order to avoid labelling primary amines in Tris. After addition of a two-molar excess of reactive NT647 dye to the respective target protein, the reaction was incubated in the dark for 30 min. After this, the unbound dye was removed using a size exclusion column as indicated by the manufacturers. Using the buffer 1X PBS pH 7.5, + 0.1% Pluronic F127 + 2% DMSO did not result in aggregation or sticking effects for HLA-A, HLA-B, MAPKAPK2, CDK2, and PKC eta, ITK, and ERα. Standard, premium and hydrophobic capillary types were tested for non-specific sticking of the proteins to the glass surface. HLA-A and HLA-B, and ERα showed sticking in standard capillaries but no sticking in premium capillaries. Hence, premium capillaries were used for the further experiments for these proteins. The other proteins remained in solution in standard capillaries. PDE4B showed aggregation in all conditions and hence was not tested further. The LED power was set to 10-25 % to obtain optimal signal intensities. The laser power was identified being optimal at 40% or 80% laser power. Changes in amplitudes between the lower and upper binding curve plateau of more than 4 units and a signal to noise ratio of above 6 were considered significant for binding events. Each experiment had 2 replicates.

## Results and Discussion

### Structural Signatures

We constructed the structural signature from the positive dataset (extracted from [11]) using sequence-independent structure alignment, hierarchical clustering and a probabilistic scoring function (see methods for details, Figure 1 (a-c)). This method has been successful in representing the binding pocket signature of eleven metabolites (drugs) in our earlier study [38]. However, in this study we found that the positive structural signature alone is not sufficient in the case of kinase inhibitors to distinguish targets from non-targets. Indeed, for these drugs, we obtained an average sensitivity and specificity of 31(±34)% and 78(±25)% respectively using the cut-off of 0.85 specified in our previous study (Table 1, Supplementary Table 2). This large standard deviation suggests that the algorithm’s performance based on using only a positive signature is not very reliable when the targets and non-targets share significant similarity. This might be due to a combination of reasons, i) the ATP binding pocket is structurally conserved across the kinase domains, ii) the orientation of the DFG motif differs across the kinase domains, and iii) there are subtle changes in the binding interaction of the kinase inhibitors with the kinase domains [42]. To resolve these issues, we built a structural signature from the negative dataset (also extracted from [11]) dubbed “negative signature” using the same procedure as the positive dataset (Figure 1 (d-f)). The pocket that was most similar to the bound pocket from each structure was used to construct the negative signature. This is similar to the well-established practice of using near-native decoys to improve the docking based scoring functions [43]. The number of structures used to make the positive and negative signatures and the preservation ratio for each signature is given in Supplementary Table 3. A protein is considered a target only when one of its top three largest pockets has a better (lower) score (as defined in the methods section) after aligning with the positive structural signature as compared to the score of the same pocket aligning with the negative structural signature (Score_positive_ – Score_negative_ < 0). Combining the positive and negative structural signatures helped improve the performance of the methodology significantly across all the kinase inhibitors (Table 1). Using a data set of all kinases with experimentally determined structures and 5 FDA-approved kinase inhibitors, we can predict the known targets with 52% sensitivity (ranging between 42% - 68%) and 55% specificity (ranging between 50% - 59%) in 5-fold cross-validation tests. To evaluate the effect of sequence similarity in the dataset we constructed a baseline model using sequence similarity (see Methods). Using this baseline sequence model, we could only assign 47% of the proteins to either the interacting or non-interacting cluster with a sensitivity and specificity of 36(±8) % and 33(±9) % respectively.

**Table 3.** Predicted interaction of Kinase Inhibitors with ERα. Interaction with ERα. sunitinib, dasatinib and pazopanib were among the top predicted targets using our method. The Score_positive_, Score_negative_, Score_positive_ – Score_negative_, docking score, significance measure (random chance to obtain a better docking score) and the experimental binding affinity are shown here.

| Drug | Score Pos | Score Neg | Score Pos-Neg | Docking Score | Random Chance | Binding Affinity |
| --- | --- | --- | --- | --- | --- | --- |
| sunitinib | 0.52 | 0.74 | -0.22 | -154.7 | 7% | 14.7 ± 5.7nM |
| dasatinib | 0.51 | 0.75 | -0.24 | -52.83 | 8% | 1.2 ± 0.5uM |
| pazopanib | 0.57 | 0.76 | -0.19 | -43.23 | 3% | 3.2 ± 1.0uM |

In our previous study, we constructed a synthetic negative dataset (as negative results are usually not published) to assess the specificity of our methodology [38]. Here, we found that using a synthetic negative dataset can severely over-estimate specificity. For example, for sorafenib using the positive signature alone, we found the specificity on the negative dataset constructed by [38] to be 65%, while on the negative dataset based on experimental results of [11] we found the specificity to be 33%. In the case of kinase inhibitors, experimental kinome profiling companies like DiscoverX have explored kinase inhibitor interactions extensively [11]. According to our data set, the 5 kinase inhibitors chosen in this study interact with 26% (on average) of the kinases and the rest are considered as true negatives. Any novel interaction discovered from these negative examples are therefore non-trivial as the model is trained to treat them as negative. The probability of discovering 3 novel interactions amongst the top 10 predictions can be determined using combinatorics and is very small (<1%).

### Identifying new targets

In order to identify new drug targets, we extracted the top 3 pockets (largest volume) of structures deposited in the Protein Data Bank (PDB) using CASTp [39]. We then aligned the structural signatures (both positive and negative) of each drug with these pockets. Similar to self-validation tests, a protein is considered a potential target only when one of the pockets has Score_positive_ – Score_negative_ < 0. We then predicted the strength of the interaction between the drug and the potential target using the flexible docking option of SwissDock [44]. To address the relatively high false positive rate and the imperfections in the scoring functions associated with docking methods, we provide a significance measure for these binding scores. The significance measure is calculated by comparing the binding scores obtained for potential targets of a drug with the binding scores of 100 random protein structures with the respective drug. The size of the random sample can be increased for improved statistical significance at the cost of significant computational time. The random structures are sampled from a list of protein structures that have less than 60% identity with the positive and the negative data set. The significance measure enables us to identify more promiscuous compounds such as gefitinib, where even the targets with most favorable docking score have an unfavorable significance measure (meaning that the compound is unusually sticky and is interacting with many proteins with high probability). Therefore, we exclude gefitinib from our study. Table 2 gives the top 10 predicted targets of sorafenib after redundancy reduction using the PISCES webserver [45]. The top 10 predicted targets for the rest of the kinase inhibitors are given in Supplementary Tables 4-7. Our ranking consists of two steps. The first step requires identifying all protein targets for which Score_positive_ – Score_negative_ < 0. In the second step, we sort in ascending order with respect to the docking score.

**Table 4.**
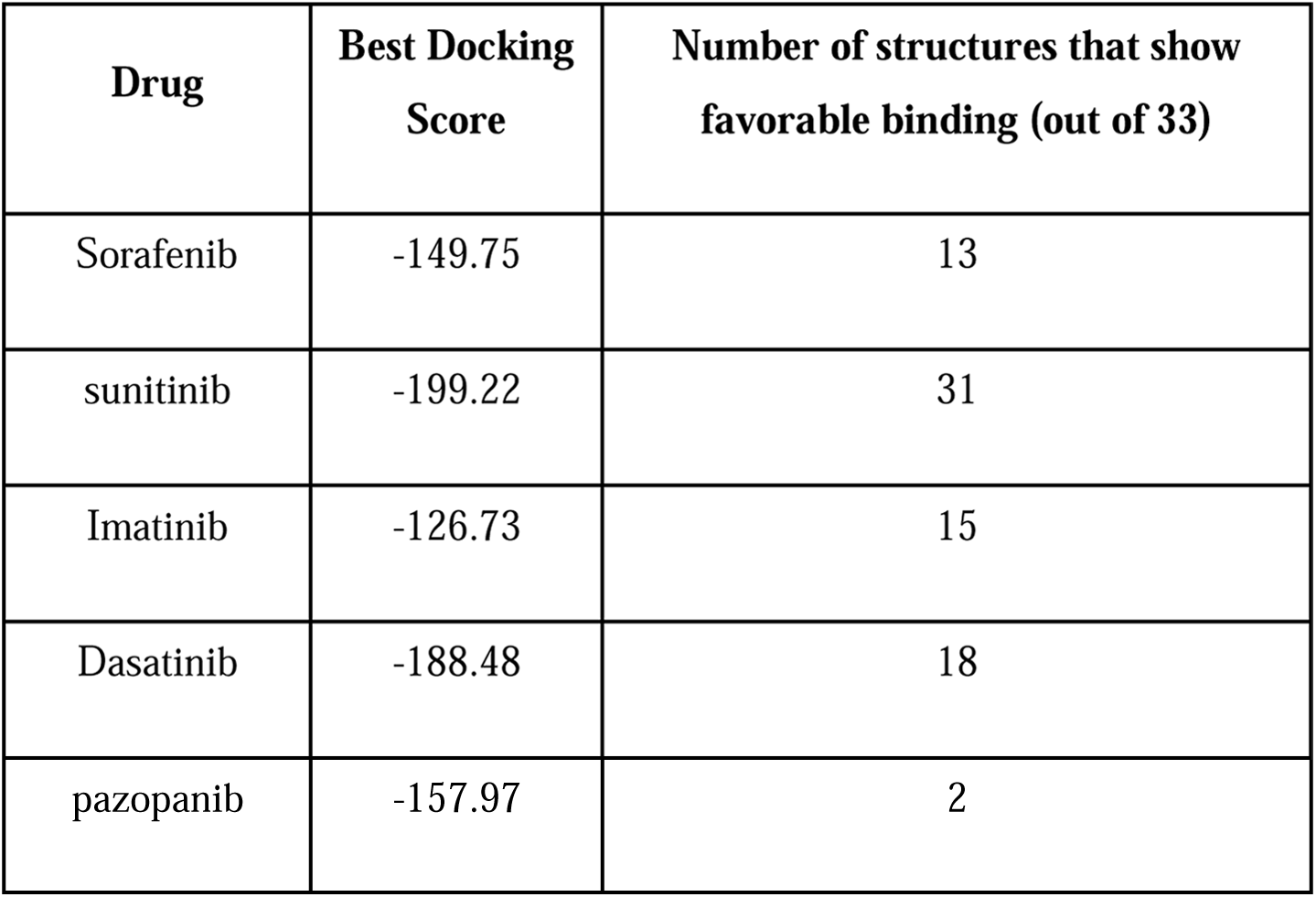
Predicted interaction of Kinase Inhibitors with MHC Class I proteins. All kinase inhibitors in this study except pazopanib were predicted to directly interact with a significant number of MHC Class I proteins (HLA alleles).

| **Drug** | **Best Docking Score** | **Number of structures that show favorable binding (out of 33)** |
| --- | --- | --- |
| Sorafenib | -149.75 | 13 |
| sunitinib | -199.22 | 31 |
| Imatinib | -126.73 | 15 |
| Dasatinib | -188.48 | 18 |
| pazopanib | -157.97 | 2 |

### Experimental Validation

To provide an experimental validation of iDTPnd, we first chose 5 predicted targets (PDE4B, HLA-A, HLA-B, MAPKAPK2, PKC η) of sorafenib. We used microscale thermophoresis (MST) experiments to test the predicted interaction *in vitro* (see methods for details). The choice of which predicted target to test resulted from a combination of the target’s ranking in our results, availability and cost of the purified protein. MST showed that PKC η and MAPKAPK2 interacted with sorafenib with a dissociation constant (Kd) of 1.1±0.4 μM and 3.7±0.1 μM respectively (Figure 2). Affinities to the primary targets of sorafenib are comparable (most of which are reported to be within 100 nM and 1 μM [11]), indicating that these interactions with PKC η and MAPKAPK2 might be pharmacologically relevant and hence valuable for medicinal chemists. Similarly, our results suggest that sorafenib interacts with HLA-A and HLA-B with Kd values between 300-600 μM. It is speculated, but plausible, that due to intracellular accumulation of sorafenib in some cell types and the wide diversity of HLA isotypes, this weak *in vitro* interaction could be clinically relevant [46,47]. For example, severe drug-specific adverse effects of sorafenib are reported to be associated with HLA-A24 sub-type of HLA-A proteins [48]. Moreover, the immune system is compromised while taking sorafenib and flu vaccination is not recommended during this period [49]. Finally, HLA-B has been shown to directly interact with a small molecule drug [50]. PDE4B was the top predicted target in our study but it showed aggregation under all tested conditions and hence the results were inconclusive for this protein. Next, we tested the predicted interaction between imatinib and ITK to see if the methodology worked for kinase inhibitors other than sorafenib. We chose to validate ITK-imatinib interaction as it was the top ranked prediction (most favorable docking score) among all predicted interactions in our results. In MST, imatinib interacted with ITK with a Kd of 550 ± 120 nM. Confirmation of this interaction suggested our method detects true, previously unrecognized, direct physical interactions and so we proceeded to evaluate predicted interactions for additional kinase inhibitors in our dataset.

**Figure 2.**
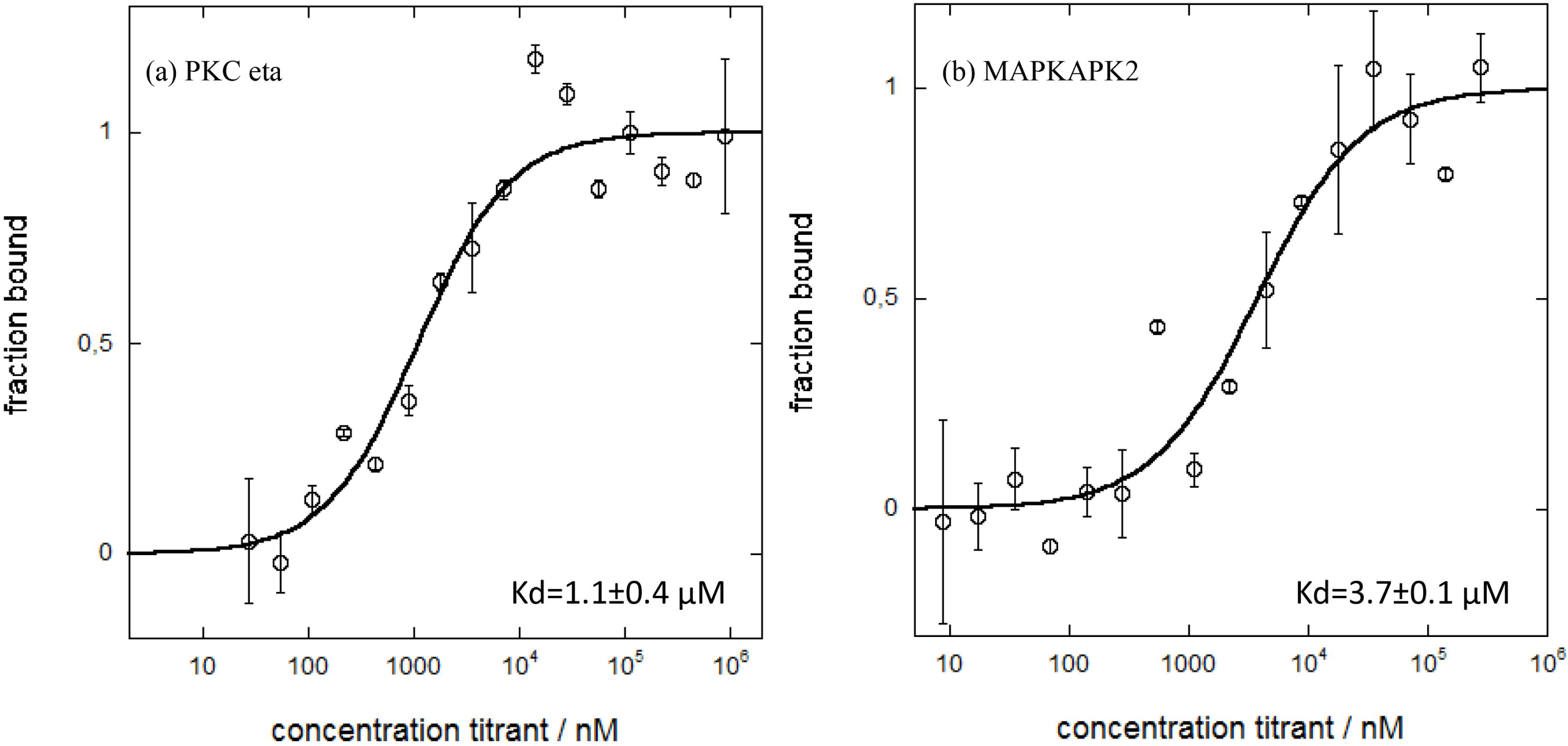
Interaction of Sorafenib with PKC η and MAPKAPK2. The predicted interaction of sorafenib with PKC η and MAPKAPK2 proteins was experimentally verified through MST experiments. MST-derived binding curve of NT647-labelled PKCη and MAPKAPK2 to sorafenib, plotted as a function of sorafenib concentration. Data are means ± S.D., n = 2.

#### Estrogen Receptor (ER)α

ERα is a nuclear receptor that is activated by estrogen and is important for hormone/DNA binding and transcription activation [51]. The role of ERα in breast cancer is well documented with nearly 70% of newly diagnosed breast cancers being ER positive (cancer cells grow in response to the hormone estrogen) [52]. ERα is one of the primary targets of tamoxifen, an FDA approved drug for breast cancer treatment [53]. ERα was ranked 3^rd^, 9^th^ and 10^th^ among the predicted targets for sunitinib, pazopanib and Dasatinib respectively in our results. We used MST to test our predictions *in vitro*. We also included sorafenib and imatinib in our experiments to test our false negative rate.

Sunitinib, dasatinib and pazopanib were found to interact with ERα with a Kd of 14.7±5.7 nM, 1.2±0.5 μM, 3.2±1.0 μM respectively (Figure 3). While sorafenib did not interact with ERα as predicted at detectable levels in our setup, we found that imatinib bound to ERα with a Kd of 335±114 nM even though ERα was not predicted as a target for imatinib in our results suggesting that iDTPnd does have some false negatives. In support, a recent case study reported response of the patient’s ER+ HER2-breast cancer tumors to pazopanib after the tumors had developed resistance to endocrine therapy [54]. Although the focus of the study was on the mechanism-independent relationship between pazopanib and fibroblast growth factor receptors, and amplified FGFR1 in the tumor. The direct interaction between pazopanib and ERα might have contributed to this clinical response. Another study shows that dasatinib can block ERα facilitated extranuclear actions that lead to metastasis [55]. This regulation can be due to the direct interaction between dasatinib and ERα. Sunitinib has also been reported to inhibit tumor growth in breast cancer cells [56]. Further studies are required to comprehensively understand the pharmaceutical effects of these interactions.

**Figure 3.**
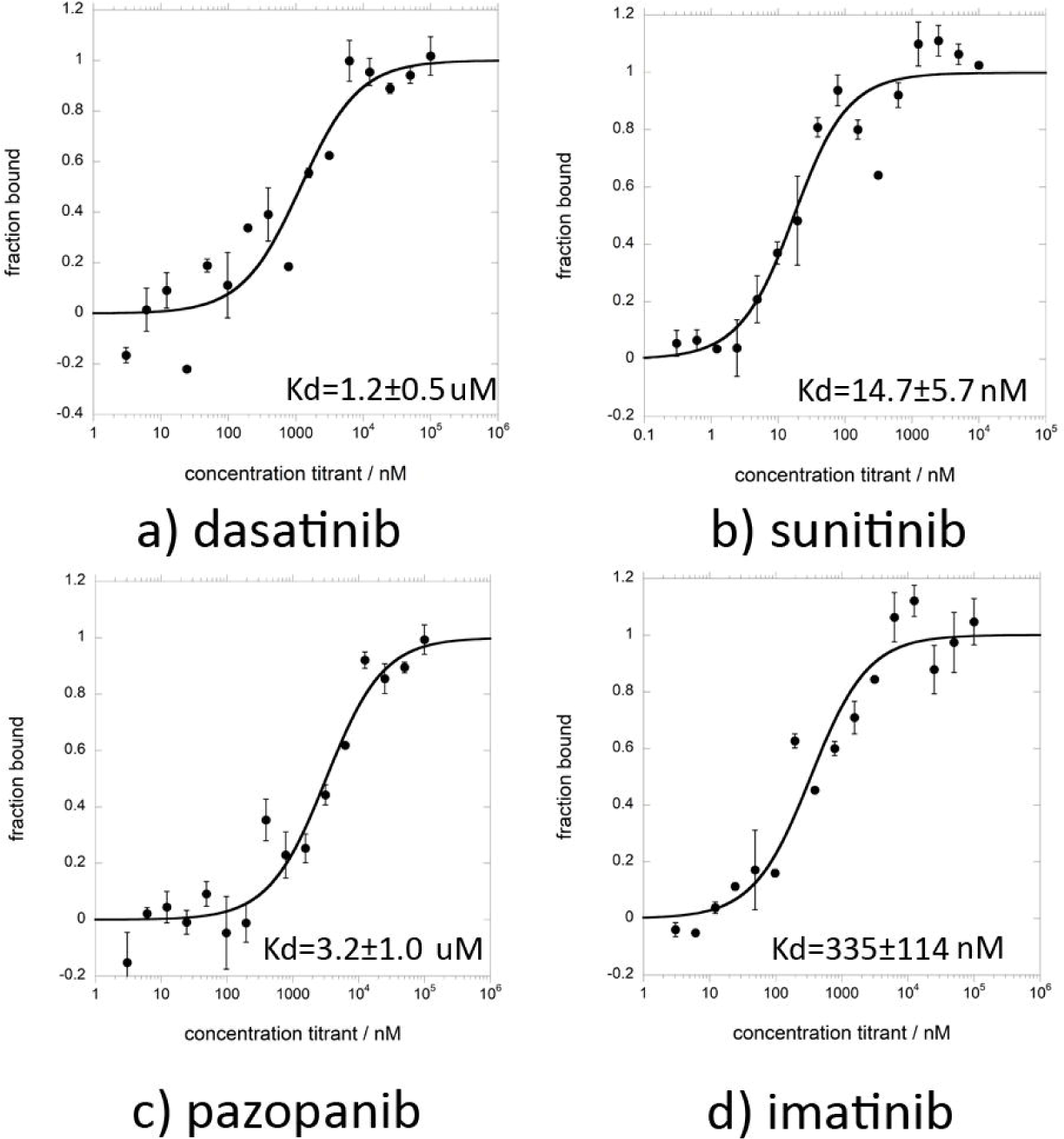
Interaction of Kinase Inhibitors with ERα. Sunitinib, dasatinib, pazopanib and Imatinib were found to interact with ERα with a Kd of 14.7±5.7 nM, 1.2±0.5 μM, 3.2±1.0 μM and 335±114 nM respectively. MST-derived binding curve of NT647-labelled ERα to ligands, plotted as a function of ligand concentration. Data are means ± S.D., n = 2.

#### Cyclin-dependent kinase 2 (CDK2)

Cyclin-dependent kinases (CDKs) perform important roles in cell division cycle, transcription, differentiation, neuronal functions and apoptosis [57]. Specifically, CDK2 has been implicated in prostate cancer, non-small cell cancer, breast cancer [58–60]. Several CDK2 inhibitors have been developed to check aberrant CDK2 activity. Sorafenib has been shown to interact with CDK2 [11]. CDK2 also appeared as one of the top targets of dasatinib and imatinib in our *in silico* prediction. In our next round of MST experiments, we also included sorafenib as the positive control and two other kinase inhibitors (sunitinib and pazopanib) in our experiments to test our false negative rate.

Dasatinib, imatinib and sorafenib were found to interact with CDK2 with a Kd of 2.2±0.9 μM, 6.6±2.9 μM, 9.1±2.7 μM respectively (Figure 4). We found that pazopanib also interacts with CDK2 with a Kd of 4.7±1.4 μM even though CDK2 was not predicted as a target for pazopanib in our results (Figure 4). While sunitinib did not interact with CDK2 as predicted at detectable levels by iDTPnd. The interaction between CDK2 and dasatinib is indirectly supported by a previous study that shows selective modulation of CDK2 by dasatanib [61]. Our results indicate that this modulation is a direct result of the interaction between CDK2 and dasatinib.

**Figure 4.**
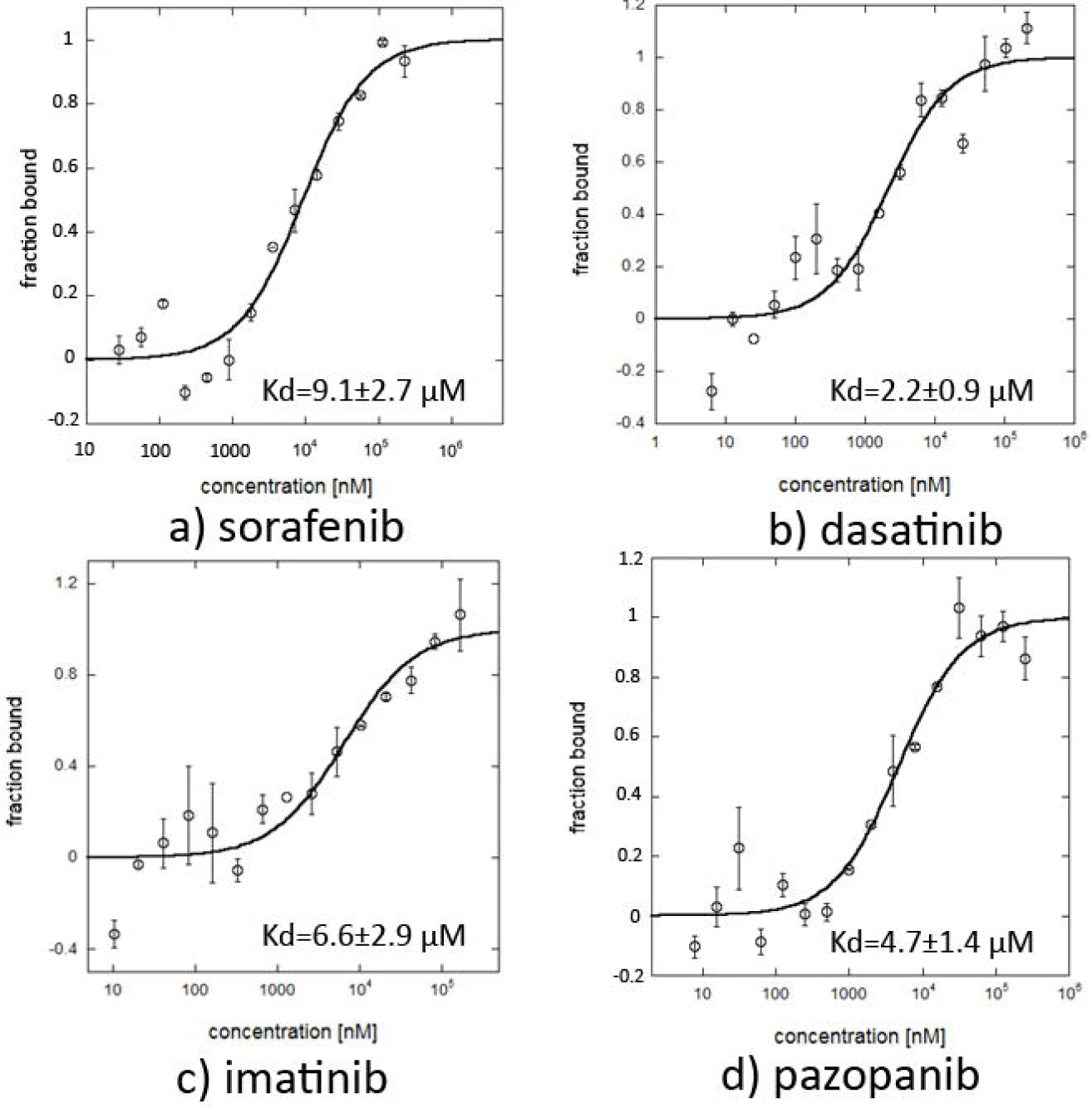
Interaction of Kinase Inhibitors with CDK2. Dasatinib, Imatinib, sorafenib and pazopanib were found to interact with CDK2 with a Kd of of 2.2±0.9 μM, 6.6±2.9 μM, 9.1±2.7 μM and 4.7±1.4 μM respectively. MST-derived binding curve of NT647-labelled CDK2 to ligand, plotted as a function of ligand concentration. Data are means ± S.D., n = 2.

#### MHC Class I proteins

We observed that MHC Class I (HLA-A/HLA-B) proteins were predicted as potential targets for all Kinase Inhibitors used in this study except pazopanib. The cell surface of all nucleated cells contain MHC Class I proteins in jawed vertebrates [62]. They bind peptides (formed due to degradation of cytosolic proteins) and display them to the cytotoxic T cells. Cytotoxic T cells bind the presented peptide and on recognition of an infected state, initiate an immune response. Peptide binding to the MHC Class I proteins is the most selective step in the antigen presentation pathway. As of August 2016, there were 33 (sequence similarity < 99%) experimentally resolved structures available for different alleles of MHC Class I proteins. To explore the kinase inhibitor – MHC Class I interactions further we performed the flexible docking of all the kinase inhibitors in this study with each of the 33 structures. The dominant interaction (determined using flexible docking) between the kinase inhibitors and the MHC Class I proteins exists in the peptide binding region (Figure 5), which is one of the two pockets identified by the structural signatures. This interaction in the binding region is significant as it might change the peptides being presented to cytotoxic T cells as in the case of abacavir, which is FDA-approved for HIV treatment [63]. The second pocket identified is located between the two chains of the MHC Class I proteins. Our results suggest that with the exception of pazopanib all kinase inhibitors tested in this study directly interact with many HLA alleles (13-31 out of 33) (Table 3). This interaction might compete with the peptides being presented to cytotoxic T cells. The direct interaction between the kinase inhibitors and MHC Class I proteins might initiate an immune response that is responsible for the observed side effects.

**Figure 5.**
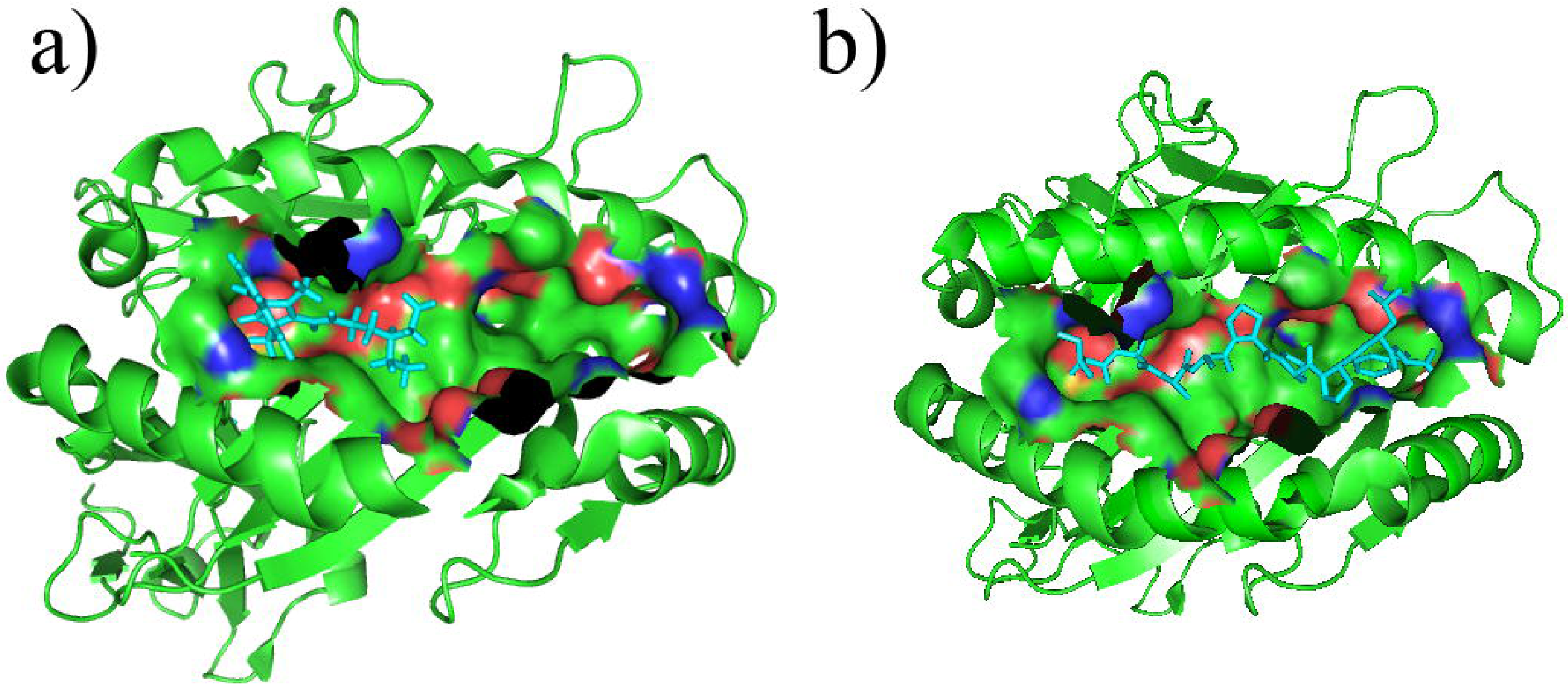
Ligand Binding Pocket of MHC Class I proteins. The extracellular ligand binding grove of MHC Class I proteins, taken from PDB 3bxn is shown as green ribbon. The ligand binding pocket identified by iDTPnd, where apolar atoms are shown in green, negatively charged atoms in blue and positively charged atoms in red. a): Sorafenib bound in the pocket (as a result of the docking). b) peptide bound in the pocket (in the crystal structure).

#### Cytochrome p450 (CYP)

CYPs are the most important enzymes involved in drug metabolism. They account for about 75% of the total metabolism. Most drugs are deactivated by CYPs, either directly or indirectly [64]. Drug metabolism by CYPs is a key reason of adverse drug interactions, as altered CYP enzyme activity can affect the metabolism and removal of drugs from the body [64]. In our results, CYP2E1 and CYP2A6 are among the top predicted targets of dasatinib and CYP1A2 is among the top predicted targets of pazopanib. As CYPs play an important role in the drug metabolism, FDA tests these interactions before approving a drug. We found that the predicted interactions were indeed reported in the FDA Orange Books (Table 5) [65,66]. Our methodology might be a good platform for drug companies to test the interaction of novel drugs with different CYP enzymes.

**Table 5.** Kinase Inhibitor – CYP450 interaction Validation. Kinase inhibitors predicted to interact with CYP450 by iDTPnd. These interactions are also reported in the FDA Orange Books.

| Drug | Sub type | Docking Score | IC50 |
| --- | --- | --- | --- |
| Dasatinib | Cytochrome P450 2E1 | -194.61 | >50μM |
| Dasatinib | Cytochrome P450 2A6 | -100.57 | 35μM |
| pazopanib | Cytochrome P450 1A2 | -105.97 | 16μM |

### Comparison with previous methods

Due to the importance of drug-protein interactions, several computational studies have addressed the problem of identifying novel targets of drugs from different angles. However, most of these studies do not benchmark their performance on known targets and use different datasets, thus making it hard to compare among these studies. The studies that do report have relatively low precision values (29%, 30% and 49%, respectively) [17,19,21]. Moreover, it is well established in literature that a dataset of confirmed negative relationships (not the negative dataset generated by randomly sampled drugs and potential targets) are pertinent to the improvement of drug target predictions [67,68]. Cichonska et al. have used machine learning methods to predict the binding affinities of kinase inhibitors to the kinome [33]. Although the authors report some success, it is not obvious to choose the kernels and regularizing parameters for applying the methodology to new drugs. Moreover, it is surprising that 3D features for both drugs and targets do not improve the performance of the methodology. Here we show that the weakly conserved features of the 3D drug binding site are sufficient to predict the binding affinity of the kinase inhibitors to the proteins whose 3D structures have been resolved. Merget et al. used machine learning to develop a kinase profiling method. Although they reported considerable success (area under the curve > 0.7), the authors did not experimentally validate new predictions [30]. Al-Ali et al. combined cell-based screening with machine learning to correlate the kinase inhibition profile to neurite growth [69]. This investigation has relative specificity for neuronal cells and requires more intensive experimental, cell-based screening.

Herein, we propose “iDTPnd”: a computational method for large-scale discovery of novel targets of known drugs. Our method has the following advantages: (i) it incorporates a negative structural image into the probabilistic scoring function increasing the sensitivity (cut-off = 0.85 as mention in [38]) from 31% to 52%. (ii) It provides a docking-based interaction score and a measure of the statistical significance of the interaction score enabling us to identify especially promiscuous small molecules like gefitinib. (iii) The performance of the scoring function is supported by *in vitro* binding experiments that validated 10 predicted interactions. Moreover, we have also compared our model with a recently published studies of Zhou *et al.* [17] and Luo *et al.* [34]. We analyzed the predicted targets of kinase inhibitors in our dataset by Zhou *et al.*’s webserver “Dr. PRODIS” [17] and the predicted targets of DTINet [34]. It is important to note that we cannot ensure training/testing data split on these tools and hence the reported results can be considered as a best case scenario. Dr. Prodis predicted 7469, 6483, 6263 and 7394 targets for Sorafenib, Imatinib, Dasatinib and Sunitinib respectively and did not give any results for Pazopanib. Similarly, DTINet predicted 2966 targets for Sorafenib, Imatinib, Dasatinib and Sunitinib respectively but did not give any results for Pazopanib. We analyzed the Top50 and Top200 targets for each drug from Dr. PRODIS and DTINet for the known proteins that contain a kinase domain and interact with the respective drugs (Table 6). These results represent the apparent sensitivity of Dr. PRODIS and DTINet, which is much lower than the sensitivity of iDTPnd.

**Table 6.** Performance of Dr. Prodis and DTINet. The average sensitivity by analyzing the top50 (top200) predictions of Dr. Prodis and DTINet is 14%(16%) and 17%(4.3%) respectively. This much lower the sensitivity of iDTPnd.

|  | Dr. PRODIS |  | DTINet |  |
| --- | --- | --- | --- | --- |
| Drug | Top50 | Top200 | Top50 | Top200 |
| sorafenib | 18% | 7.5% | 14% | 3.5% |
| dasatinib | 10% | 21.5% | 20% | 5% |
| imatinib | 14% | 27.5% | 18% | 4.5% |
| sunitinib | 14% | 8% | 16% | 4% |
| pazopanib | --- | --- | --- | --- |
| Average | 14% | 16% | 17% | 4.3% |

### Application to allosteric binding sites

ATP binding site has conserved features across most kinase domains, several kinase inhibitors interact with the human kinome broadly and are not very selective. However, type IV inhibitors bind to allosteric sites that are topologically and spatially distinct from conserved ATP-binding sites. It is natural to extend our methodology to allosteric biding sites. However, allosteric binding sites are significantly different than non-allosteric binding sites in terms of shape and residue conservation [70]. The construction of the structural signature requires at least 50% conservation for each position in the signature. Therefore, we plan on exploring the application of iDTPnd in detail on allosteric binding sites in future studies.

## Conclusion

We have developed a computational model “iDTPnd” to discover the novel targets of known drugs. For the five kinase inhibitors in our dataset, we can identify the known targets with 52% sensitivity and 55% specificity. The predictive capability of the methodology was supported by the validation of top predicted targets using *in vitro* binding experiments. First, we showed that 4 of the top 10 predicted targets of sorafenib were binders. PKC eta and MAPKAPK2 had Kd similar to the primary targets of sorafenib, it is therefore possible that these interactions can be exploited in various cancer treatments. Similarly, the interaction between sorafenib and MHC Class I proteins might play currently unexplored roles in immune response to kinase inhibitors. Previously abacavir, an HIV protease inhibitor, has been shown to alter the peptide binding preference of MHC Class I molecules. It is probable that same might be true for several kinase inhibitors.

Second, we verified kinase inhibitor interaction with two proteins (ERα and CDK2) that appeared in the top 10 predicted target list of more than one kinase inhibitors. In both cases our predicted interactions were verified by *in vitro* experiments. Beyond validating our predictions, experimental results also suggest that our method can serve as a platform for kinase inhibitor combination studies. The experimental validation shows that our false positive rate is very low compared with other studies. The false negative rate can be improved in future studies by incorporating structure independent information like expression data from GTEx and ENCODE projects. Our methodology is generic and can be used broadly for all types of small molecule drugs for which sufficient (~30) 3D structures of known targets have been solved.

## Data Availability

The code for constructing the structural signature is available at https://sfb.kaust.edu.sa/Documents/iDTP.zip

## Author Contributions

HN designed research; HN performed research; CK and TS performed the experimental validation; HN, STA, XG and MLM analyzed data; HN, STA, XG and MLM wrote the paper.

## Acknowledgements

The authors thank Dr. Aly Azeem Khan for helpful discussions. This work has been supported by the Toyota Technological Institute at Chicago, King Abdullah University of Science and Technology and a grant to establish Precision Medicine Lab under the umbrella of National Center in Big Data & Cloud Computing from the Higher Education of Pakistan. The research by STA reported in this publication was supported by funding from King Abdullah University of Science and Technology (KAUST), Office of Sponsored Research (OSR), under award number FCC/1/1976-25.

## Supplementary Information

**Supplementary Figure 1 Flow chart of the methodology**

**Supplementary Table 1 Dataset**

The pdb structures used to make the respective positive structural signatures (pockets extracted from the first chain) and negative signatures (pocket most similar to the bound pocket).

**Supplementary Table 2 Performance using positive signature alone**

The positive structural signature alone is not sufficient in the case of kinase inhibitors to distinguish targets from non-targets as using different cut-offs the sensitivity or specificity becomes unacceptable. Values less than 60% are highlighted.

**Supplementary Table 3 Positive and Negative Signature**

The number of structures used to make the positive and negative signatures and the redundancy cut-off for each signature.

**Supplementary Table 4 Predicted Targets of Dasatinib**

Top 10 predicted targets of Dasatinib, their PDB id, Score_positive_, Score_negative_, Score_positive_ – Score_negative_, docking score and the significance measure (random chance to obtain a better docking score).

**Supplementary Table 5 Predicted Targets of Imatinib**

Top 10 predicted targets of Imatinib, their PDB id, Score_positive_, Score_negative_, Score_positive_ – Score_negative_, docking score and the significance measure (random chance to obtain a better docking score).

**Supplementary Table 6 Predicted Targets of Sunitinib**

Top 10 predicted targets of Sunitinib, their PDB id, Score_positive_, Score_negative_, Score_positive_ – Score_negative_, docking score and the significance measure (random chance to obtain a better docking score).

**Supplementary Table 7 Predicted Targets of Pazopanib**

Top 10 predicted targets of Pazopanib, their PDB id, Score_positive_, Score_negative_, Score_positive_ – Score_negative_, docking score and the significance measure (random chance to obtain a better docking score).

## References

[1] Johnson LN, Lewis RJ. Structural basis for control by phosphorylation. Chem. Rev. 2001; 101: 2209–2242.

[2] Adams JA. Kinetic and catalytic mechanisms of protein kinases. Chem. Rev. 2001; 101: 2271–2290.

[3] UniProt Consortium. UniProt: a hub for protein information. Nucleic Acids Res. 2015; 43 (Database issue): D204–12.

[4] Knighton DR, Zheng JH, Ten Eyck LF, Ashford VA, Xuong NH, Taylor SS, et al. Crystal structure of the catalytic subunit of cyclic adenosine monophosphate-dependent protein kinase. Science. 1991; 253: 407–414.

[5] Huang M, Shen A, Ding J, Geng M. Molecularly targeted cancer therapy: some lessons from the past decade. Trends Pharmacol. Sci. 2014; 35: 41–50.

[6] Ma WW, Adjei AA. Novel agents on the horizon for cancer therapy. CA Cancer J. Clin. 2009; 59: 111–137.

[7] Sun C, Bernards R. Feedback and redundancy in receptor tyrosine kinase signaling: relevance to cancer therapies. Trends Biochem. Sci. 2014; 39: 465–474.

[8] Noble ME, Endicott JA, Johnson LN. Protein kinase inhibitors: insights into drug design from structure. Science. 2004; 303: 1800–1805.

[9] Norman RA, Toader D, Ferguson AD. Structural approaches to obtain kinase selectivity. Trends Pharmacol. Sci. 2012; 33: 273–278.

[10] Karaman MW, Herrgard S, Treiber DK, Gallant P, Atteridge CE, Campbell BT, et al. A quantitative analysis of kinase inhibitor selectivity. Nat Biotechnol. 2008 Jan; 26(1):127–32.

[11] Davis MI, Hunt JP, Herrgard S, Ciceri P, Wodicka LM, Pallares G, et al. Comprehensive analysis of kinase inhibitor selectivity. Nat. Biotechnol. 2011; 29: 1046–1051.

[12] Arrowsmith J. Trial watch: phase III and submission failures: 2007–2010. Nat. Rev. Drug Discov. 2011; 10: 87.

[13] Arrowsmith J. Trial watch: Phase II failures: 2008–2010. Nat. Rev. Drug Discov. 2011; 10: 328–329.

[14] Liebler D, Guengerich F. Elucidating mechanisms of drug-induced toxicity. Nat. Rev. Drug Discov. 2005; 4(5): 410–420.

[15] Maitland ML, Ratain MJ. Terminal ballistics of kinase inhibitors: there are no magic bullets. Ann Intern Med. 2006; 145(9): 702–3.

[16] Lounkine E, Keiser MJ, Whitebread S, Mikhailov D, Hamon J, Jenkins JL, et al. Large-scale prediction and testing of drug activity on side-effect targets. Nature. 2012; 486(7403): 361–367.

[17] Zhou H, Gao M, Skolnick J. Comprehensive prediction of drug-protein interactions and side effects for the human proteome. Scientific Reports. 2015; 5:11090.

[18] Hughes JP, Rees S, Kalindjian SB, Philpott KL. Principles of early drug discovery. Br. J. Pharmacol. 2011; 162: 1239–1249.

[19] Chang R, Xie L, Xie L, Bourne P, Palsson B. Drug mechanism-independent effects predicted using structural analysis in the context of a metabolic network model. PLoS Comput. Biol. 2010; 6(9): e1000938.

[20] Engin HB, Keskin O, Nussinov R, Gursoy A. A strategy based on protein-protein interface motifs may help in identifying drug mechanism-independents. J. Chem. Inf. Model. 2012; 52: 2273–2286.

[21] Li YY, An J, Jones SJ. A computational approach to finding novel targets for existing drugs. PLoS Comput. Biol. 2011; 7: e1002139.

[22] Hwang H, Dey F, Petrey D, Honig B. Structure-based prediction of ligand-protein interactions on a genome-wide scale. Proc. Natl. Acad. Sci. U S A. 2017; 114 (52): 13685–13690.

[23] Emig D, Ivliev A, Pustovalova O, Lancashire L, Bureeva S, Nikolsky Y, et al. Drug target prediction and repositioning using an integrated network-based approach. PLoS One. 2013; 8: e60618.

[24] Hu G, Agarwal P. Human disease-drug network based on genomic expression profiles. PLoS One. 2009; 4(8): e6536.

[25] Iorio F, Bosotti R, Scacheri E, Belcastro V, Mithbaokar P, Ferriero R, et al. Discovery of drug mode of action and drug repositioning from transcriptional responses. Proc. Natl. Acad. Sci. U S A. 2010; 107(33): 14621–14626.

[26] Suthram S, Dudley J, Chiang A, Chen R, Hastie T, Butte A. Network-based elucidation of human disease similarities reveals common functional modules enriched for pluripotent drug targets. PLoS Comput. Biol. 2010; 6(2): e1000662.

[27] Wei G, Twomey D, Lamb J, Schlis K, Agarwal J, Stam R, et al. Gene expression-based chemical genomics identifies rapamycin as a modulator of MCL1 and glucocorticoid resistance. Cancer Cell. 2006; 10(4): 331–342.

[28] Keiser M, Setola V, Irwin J, Laggner C, Abbas A, Hufeisen S, et al. Predicting new molecular targets for known drugs. Nature. 2009; 462(7270): 175–181.

[29] Qu X, Gudivada R, Jegga A, Neumann E, Aronow B. Inferring novel disease indications for known drugs by semantically linking drug action and disease mechanism relationships. BMC Bioinformatics. 2009; 10 Suppl 5, S4.

[30] Merget B, Turk S, Eid S, Rippmann F, Fulle S. Profiling prediction of kinase inhibitors: toward the virtual assay. J Med. Chem. 2016; 60: 474–485.

[31] Napolitano F, Zhao Y, Moreira VM, Tagliaferri R, Kere J, D’Amato M et al. Drug repositioning: a machine-learning approach through data integration. J. Cheminform. 2013; 5: 30.

[32] Wang Z, Clark NR, Ma’ayan A. Drug-induced adverse events prediction with the LINCS L1000 data. Bioinformatics. 2016; 32(15):2338–45.

[33] Cichonska A, Ravikumar B, Parri E, Timonen S, Pahikkala T, Airola A, et al. Computational-experimental approach to drug-target interaction mapping: A case study on kinase inhibitors. PLOS Comput Biol. 2017; 13(8): e1005678.

[34] Luo Y, Zhao X, Zhou J, Yang J, Zhang Y, Kuang W, et al. A network integration approach for drug-target interaction prediction and computational drug repositioning from heterogeneous information. Nat Commun, 2017; 8, 1:573.

[35] Wan F, Hong L, Xiao A, Jiang T, Zeng J. NeoDTI: neural integration of neighbor information from a heterogeneous network for discovering new drug-target interactions. Bioinformatics, 2019; 35, 1:104–111.

[36] Campillos M, Kuhn M, Gavin A, Jensen L, Bork P. Drug target identification using side-effect similarity. Science. 2008, 321(5886), 263–266.

[37] Sanseau P, Agarwal P, Barnes M, Pastinen T, Richards J, Cardon L, et al. Use of genome-wide association studies for drug repositioning. Nat. Biotechnol. 2012, 30(4), 317–320.

[38] Naveed H, Hameed US, Harrus D, Bourguet W, Arold ST, Gao X. An integrated structure- and system-based framework to identify new targets of metabolites and known drugs. Bioinformatics. 2015, 31(24), 3922–3929.

[39] Dundas J, Ouyang Z, Tseng J, Binkowski A, Turpaz Y, Liang J. CASTp: computed atlas of surface topography of proteins with structural and topo-graphical mapping of functionally annotated residues. Nucleic Acids Res. 2006, 34(Web Server issue), W116–8.

[40] X Cui, H Naveed, X Gao. Finding optimal interaction interface alignments between biological complexes. Bioinformatics. 31 (12), i133–i141.

[41] Dundas J, Adamian L, Liang J. Structural signatures of enzyme binding pockets from order-independent surface alignment: a study of metalloendopeptidase and NAD binding proteins. J Mol Biol. 2011, 406(5), 713–29.

[42] Wu P, Nielsen TE, Clausen MH. FDA-approved small-molecule kinase inhibitors. Trends in Pharmacological Sciences, 2015, 1–18.

[43] Liu S, Vakser IA. DECK: Distance and environment-dependent, coarse-grained, knowledge-based potentials for protein-protein docking. BMC Bioinformatics, 2011, 12, 128.

[44] Grosdidier A, Zoete V, Michielin O. SwissDock, a protein-small molecule docking web service based on EADock DSS. Nucleic Acids Res. 2011, 39(Web Server issue), W270–7.

[45] Wang G, Dunbrack RL Jr. PISCES: recent improvements to a PDB sequence culling server. Nucleic Acids Res. 2005, 33, W94–8.

[46] Pratz KW, Cho E, Levis MJ, Karp JE, Gore SD, McDevitt M, et al. A pharmacodynamic study of sorafenib in patients with relapsed and refractory acute leukemias. Leukemia, 2010, 24 (8), 1437–44.

[47] Hu S, Chen Z, Franke R, Orwick S, Zhao M, Rudek MA, et al. Interaction of the multikinase inhibitors sorafenib and sunitinib with solute carriers and ATP-binding cassette transporters. Clin Cancer Res., 2009, 15(19), 6062–9.

[48] Tsuchiya N, Narita S, Inoue T, Hasunuma N, Numakura K, Horikawa Y, et al. Risk factors for sorafenib-induced high-grade skin rash in Japanese patients with advanced renal cell carcinoma. Anti-Cancer Drugs 2013, 24(3), 310–314.

[49] Hipp MM, Hilf N, Walter S, Werth D, Brauer KM, Radsak MP, et al. Sorafenib, but not sunitinib, affects function of dendritic cells and induction of primary immune responses. Blood. 2008, 111(12), 5610–5620.

[50] Wei CY, Chung WH, Huang HW, Chen YT, Hung SI, et al. Direct interaction between HLA-B and carbamazepine activates T cells in patients with Stevens-Johnson syndrome. J Allergy Clin Immunol. 2012, 129(6), 1562–1569.

[51] Dahlman-Wright K, Cavailles V, Fuqua SA, Jordan VC, Katzenellenbogen JA, Korach KS, et al. International Union of Pharmacology. LXIV. Estrogen receptors. Pharmacol. Rev. 2006, 58 (4), 773–81.

[52] Felzen V, Hiebel C, Koziollek-Drechsler I, Reißig S, Wolfrum U, Kögel D, et al. Estrogen receptor α regulates non-canonical autophagy that provides stress resistance to neuroblastoma and breast cancer cells and involves BAG3 function. Cell Death & Disease, 2015, 6, e1812.

[53] Jordan VC, Brodie AM. Development and evolution of therapies targeted to the estrogen receptor for the treatment and prevention of breast cancer. Steroids. 2007, 72, 7–25.

[54] Cheng FT, Ou-Yang F, Lapke N, Tung KC, Chen YK, Chou YY, et al. Pazopanib Sensitivity in a Patient With Breast Cancer and FGFR1 Amplification. J Natl Compr Canc Netw. 2017, 15, 1456–1459.

[55] Chakravarty D, Nair SS, Santhamma B, Nair BC, Wang L, Bandyopadhyay A, et al. Extranuclear functions of ER impact invasive migration and metastasis by breast cancer cells. Cancer Research. 2010, 70(10), 4092–4101.

[56] Belali O, Ansari M, Korashy H. Sunitinib induces growth inhibition and apoptosis in breast cancer MDA-MB-231 cells through foxo3a signaling pathway. FASEB J. 2015, 29, 619.3.

[57] Gray N, Detivaud L, Doerig C, Meijer L. ATP-site directed inhibitors of cyclin-dependent kinases. Curr. Med. Chem. 1999, 6, 859–875.

[58] Flores O, Wang Z, Knudsen KE, Burnstein KL. Nuclear targeting of cyclin-dependent kinase 2 reveals essential roles of cyclin-dependent kinase 2 localization and cyclin E in vitamin D-mediated growth inhibition. Endocrinology. 2010, 151, 896–908.

[59] Kawana H, Tamaru J-I, Tanaka T, Hirai A, Saito Y, Kitagawa M, et al. Role of p27Kip1 and cyclin-dependent kinase 2 in the proliferation of non-small cell lung cancer. Am. J. Pathol. 1998, 153, 505–513.

[60] Ali S, Heathcote DA, Kroll SH, Jogalekar AS, Scheiper B, Patel H, et al. The development of a selective cyclin-dependent kinase inhibitor that shows antitumor activity. Cancer Res. 2009, 69, 6208–6215.

[61] Nunoda K, Tauchi T, Takaku T, Okabe S, Akahane D, Sashida G, et al. Identification and functional signature of genes regulated by structurally different ABL kinase inhibitors. Oncogene, 2007, 26, 4179–4188.

[62] Hewitt EW. The MHC class I antigen presentation pathway: strategies for viral immune evasion. Immunology. 2003, 110 (2), 163–169.

[63] Ostrov DA, Grant BJ, Pompeu YA, Sidney J, Harndahl M, Southwood S, et al. Drug hypersensitivity caused by alteration of the MHC-presented self-peptide repertoire. Proc Natl Acad Sci U S A. 2012, 109(25), 9959–64.

[64] Zanger UM, Schwab M. Cytochrome P450 enzymes in drug metabolism: regulation of gene expression, enzyme activities, and impact of genetic variation. Pharmacol Ther. 2013, 138(1), 103–41.

[65] Clinical Pharmacology and Biopharmaceutics Review(s), Application Number: 22-465. FDA. Submission Date: 19 December 2008.

[66] Clinical Pharmacology and Biopharmaceutics Review(s), Application Number: 21-986 & 22-072. FDA. Submission Date: 28 December 2005.

[67] Ding H, Takigawa I, Mamitsuka H, Zhu S. Similarity-based machine learning methods for predicting drug-target interactions: a brief review. Brief. in Bioinform. 2014, 15, 734–747.

[68] Chen X, Yan CC, Zhang X, Zhang X, Dai F, Yin J, et al. Drug-target interaction prediction: databases, web servers and computational models. Brief. Bioinform. 2016, 17, 696–712.

[69] Al-Ali H, Lee DH, Danzi MC, Nassif H, Gautam P, Wennerberg K, et al. Rational polypharmacology: systematically identifying and engaging multiple drug targets to promote axon growth. ACS Chem. Biol. 2015, 10(8), 1939–1951.

[70] Lu S, He X, Ni D, Zhang J. Allosteric Modulator Discovery: From Serendipity to Structure-Based Design. J. Med. Chem. 2019.

