## Supplementary figures and images for "Identifying Novel Targets by using Drug-binding Site Signature: A Case Study of Kinase Inhibitors"

### Supplemental Figure 1

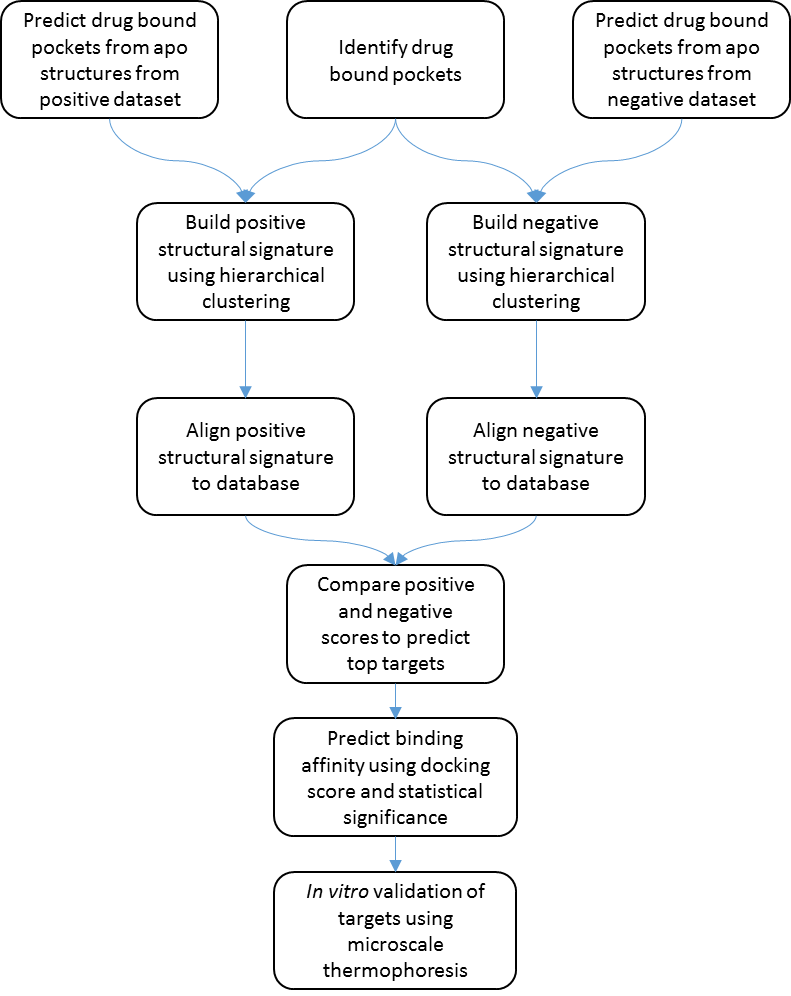

Supplementary Figure 1: Flow chart of the methodology.
