## Supplemental Table 1 for "Identifying Novel Targets by using Drug-binding Site Signature: A Case Study of Kinase Inhibitors"

| **Drugs** | **Pockets used for positive signature** | **Pockets used for negative signature** |
| --- | --- | --- |
| **sorafenib** | 2AC3 (32), 4AF3 (43), 4ASD (43), 4ASZ (41), 4BKJ (72), 3BRB (57), 4C8B (79), 4CKJ (38), 3COK (65), 4F6U (51), 3GCS (34), 3GP0 (48), 2HI4 (81), 3HMI (29), 4HVS (40), 4HW7 (31), 4I91 (84), 2J7T (31), 4JS8 (36), 3KUL (69), 4OBO (58), 2OG8 (56), 2OO8 (37), 1R9O (67), 3RGF (90), 3S95 (66), 3TBG (140), 1UA2 (39), 3UA1 (75), 1UWH (70), 4WUN (74) | 1BO1 (57), 1J1B (49), 1P4O (40), 1QCF (55), 1VZO (44), 1WAK (42), 1X8B (30), 1XJD (26), 1Z57 (47), 2A19 (28), 2BUJ (39), 2F57 (32), 2H6D (32), 2HAK (42), 2I6L (33), 2IZU (37), 2JC6 (27), 2OZO (82), 2R2P (37), 2RKU (43), 2VD5 (62), 2W4O (32), 2W5A (33), 2WNT (30), 2WQM (33), 2WTK (43), 2X7F (32), 2XIK (38), 2XRW (53), 2Y7J (31), 2Z7Q (34), 2ZV2 (37), 3A99 (32), 3ALN (36), 3BHH (33), 3CBL (41), 3CC6 (36), 3DTC (30), 3EQC (34), 3FME (30), 3FXZ (30), 3G2F (34), 3KN6 (33), 3LM5 (30), 3LXP (31), 3M2W (36), 3MI9 (50), 3MTL (40), 3OFM (54), 3OOM (35), 3P1A (35), 3PP0 (49), 3SOC (33), 3SXS (30), 3TXO (38), 3UGC (38), 3ZBF (37), 4AU8 (42), 4B6L (32), 4B9D (41), 4BF2 (28), 4BKY (47), 4CT2 (30), 4FKL (39), 4FOC (44), 4FYO (38), 4GV1 (50), 4HCU (28), 4HNI (40), 4HYI (29), 4HZR (33), 4I4E (43), 4IC7 (56), 4ITJ (37), 4J8M (31), 4JPS (16), 4L00 (34), 4LG4 (39), 4O38 (36), 4OTH (42), 4OTP (32), 4PED (69), 4PF4 (38), 4QQT (34), 4QTB (43), 4TPT (36), 4UAL (55), 4W9W (36), 4WB8 (50), 4X7Q (31) |
| **imatinib** | 4BKJ (72), 4BKY (48), 3COK (63), 4CSV (26), 3FW1 (28), 2GK9 (59), 3HEC (40), 2HI4 (81), 3HMI (29), 4HW7 (31), 3JRY (45), 3K5V (65), 3KQ0 (28), 3KUL (69), 3LXP (31), 4O38 (74), 2OIQ (67), 2PL0 (36), 4PMP (29), 1R9O (67), 1RJB (40), 1T46 (33), 3TBG (142), 3UA1 (75), 1UWJ (72), 4WKQ (37), 2WUH (22), 1XBB (38), 2XRW (56), 4YFI (53), 1Z57 (49) | 1E8Y (14), 1J1B (50), 1O43 (6), 1P4O (42), 1VZO (44), 1WAK (43), 1X8B (31), 1XJD (28), 2A19 (28), 2AC3 (33), 2BUJ (39), 2DYL (40), 2EU9 (44), 2EVA (44), 2H6D (32), 2I6L (34), 2JAM (30), 2NRU (54), 2OO8 (37), 2QOL (34), 2RKU (44), 2V62 (47), 2VWI (25), 2W4O (34), 2W5A (34), 2W96 (34), 2WEL (27), 2WQM (33), 2WTK (53), 2X4F (40), 2XIR (35), 2Y7J (32), 2YCF (45), 2ZV2 (38), 3A7I (38), 3A8X (36), 3BRB (32), 3C0I (42), 3CBL (41), 3D7T (42), 3DTC (31), 3EQC (34), 3FE3 (41), 3FME (30), 3FXZ (32), 3G2F (35), 3KVW (48), 3LFF (37), 3LM5 (32), 3M2W (37), 3MI9 (49), 3P1A (37), 3PP0 (49), 3R04 (33), 3S95 (41), 3SOC (35), 3UGC (21), 3V8S (50), 3ZBF (39), 4AAA (41), 4ASZ (41), 4B9D (41), 4BF2 (29), 4BGQ (36), 4CKJ (39), 4EUU (45), 4F6U (51), 4FKL (40), 4FOC (1), 4FR4 (49), 4GV1 (51), 4HYI (31), 4HZR (33), 4I4E (45), 4I5P (43), 4IC7 (56), 4IIR (52), 4ITJ (38), 4J8M (32), 4JPS (17), 4JS8 (36), 4KIK (84), 4KS7 (37), 4KWP (52), 4L00 (35), 4NW6 (38), 4OTH (42), 4OTP (33), 4PED (69), 4PF4 (38), 4R1V (38), 4RFZ (40), 4TWC (34), 4U3Y (44), 4UAL (56), 4USF (33), 4W9W (36), 4WB8 (52), 4WNO (35), 4X2F (37), 4YHJ (63), 4YJR (40), 4YLL (47), 4YZ9 (47), 4ZZN (46), 5ACK (29), 5AR2 (35), 5C46 (68), 5EFQ (40) |
| **dasatinib** | 1BYG (26), 4C8B (43), 3COM (44), 2EVA (44), 4FVP (37), 3GP0 (48), 2GQG (87), 2HI4 (81), 3HNG (18), 4HOK (37), 4HVS (40), 2IVT (52), 2J7T (30), 4J8M (32), 3KUL (36), 3LXL (21), 3MDY (39), 4MNE (30), 3NJP (88), 4O38 (36), 4OBO (27), 3OCT (36), 2OGV (46), 2OG8 (32), 4OTW (44), 3P1A (37), 4P5Q (38), 4PED (67), 3PM0 (70), 3PP0 (49), 3Q4U (38), 4QMS (48), 1RJB (40), 3S95 (41), 3SOC (35), 3SXR (76), 4TPT (36), 1U46 (36), 2VD5 (62), 2V62 (47), 4X2N (34), 1X8B (31), 1XBB (38), 3UA1 (75), 4UAK (57), 4WUN (43), 2WUH (22), 2Y6O (35), 2ZVA (37), 3G5D (35), 5BVW (35), 3LFA (47) | 1E8Y (14), 1J1B (50), 1P4O (4), 1U59 (46), 1UA2 (34), 1VZO (44), 1WAK (43), 1XJD (28), 1Z57 (49), 2A19 (26), 2AC3 (33), 2BUJ (39), 2CMW (40), 2DYL (40), 2EU9 (44), 2GK9 (52), 2H6D (32), 2I6L (34), 2JAM (30), 2NRU (54), 2OO8 (16), 2REI (30), 2RKU (44), 2VWI (25), 2W4O (34), 2W96 (34), 2WEL (27), 2WQM (33), 2WTK (53), 2X4F (41), 2XRW (55), 2Y7J (32), 2YCF (45), 2ZV2 (38), 3A7I (38), 3A8X (36), 3ALN (35), 3BRB (32), 3C0I (42), 3CBL (40), 3CC6 (37), 3COK (29), 3DTC (31), 3FE3 (41), 3FME (30), 3FXZ (32), 3G2F (35), 3KVW (48), 3LM5 (32), 3M2W (37), 3MI9 (50), 3MTL (42), 3R04 (34), 3TXO (38), 3V8S (50), 3WF7 (33), 3ZBF (39), 3ZDU (40), 4AAA (41), 4AF3 (36), 4AGU (34), 4ASZ (41), 4au8 (42), 4b9d (41), 4bf2 (29), 4bgq (36), 4bky (48), 4euu (45), 4f6u (51), 4fkl (40), 4foc (46), 4fr4 (49), 4gv1 (51), 4hcu (30), 4hyi (31), 4i4e (45), 4i5p (43), 4ic7 (56), 4iir (52), 4itj (38), 4jps (17), 4js8 (36), 4jsn (33), 4kik (84), 4ks7 (37), 4kwp (52), 4l00 (35), 4lg4 (41), 4nw6 (38), 4oth (42), 4otp (33), 4pf4 (38), 4qqt (35), 4r1v (38), 4twc (34), 4w9w (36), 4wb8 (52), 4wno (35), 4x7q (31), 4yhj (64), 4yll (47), 4yno (41), 4yz9 (47), 4zzn (45), 5ack (29), 5c46 (68), 5efq (40) |
| **sunitinib** | 2AC3 (33), 4AE9 (96), 4AF3 (43), 4AGD (43), 3ALO (37), 4APC (76), 4ASZ (41), 3BHH (11), 4BKJ (72), 4BKY (48), 1BO1 (6), 2BUJ (83), 3CBL (41), 1CM8 (99), 2CMW (40), 4CMT (45), 3COK (64), 3COM (85), 3DTC (31), 4FGB (42), 1FMK (44), 4FST (26), 4FVP (37), 3FZO (31), 3G0E (47), 3G2F (83), 3GP0 (48), 1H1W (32), 2H6D (32), 3HMI (29), 4HW7 (31), 3IEC (89), 1IRK (36), 4ITJ (67), 2IVT (52), 2J7T (30), 2JAM (60), 2JDO (47), 4JS8 (36), 4K00 (31), 3KN6 (67), 4KS8 (32), 1KWP (94), 3LM0 (34), 3LXL (38), 3MIY (68), 4MNE (134), 3MTL (42), 3NR9 (41), 4NY0 (121), 4O38 (74), 4OBO (58), 2OG8 (56), 4OLI (87), 2OO8 (37), 4OTD (45), 4OTP (33), 2P0C (64), 4P5Q (38), 4PF4 (38), 3Q52 (40), 3Q6U (29), 4QMZ (47), 1RJB (41), 3RNY (85), 3TI1 (44), 1U46 (84), 3U87 (6), 1UA2 (39), 3UA1 (74), 4UAK (55), 3UIU (88), 3UYS (31), 3VN9 (53), 1VZO (44), 2W4O (34), 2W5H (33), 1WAK (43), 2WQM (33), 4WSQ (78), 2WTK (138), 4WUN (74), 4X2N (34), 2X4F (79), 1X8B (31), 1XBB (38), 1XJD (28), 2XRW (56), 2Y7J (19), 1Z57 (49), 2Z7Q (35), 2ZOQ (2), 2ZV2 (38) | 3A99 (33), 4AGU (91), 5AR2 (76), 4B6L (32), 4B9D (79), 4BF2 (50), 4BGQ (36), 3BLH (78), 5C46 (92), 1CM8 (99), 3D7T (84), 1E8Z (23), 5EFQ (47), 2EU9 (42), 4EYJ (51), 4F6U (93), 4FKL (40), 4FR4 (255), 3FXZ (32), 4FYO (40), 2GK9 (157), 4GV1 (51), 5HG8 (38), 2I6L (72), 1J1B (105), 4JPS (154), 4JSP (300), 4KIK (176), 3KUL (70), 3M2W (37), 3MFS (33), 2OO8 (37), 3OOM (36), 4OTW (44), 2OZO (81), 3P1A (37), 4P2K (34), 4PED (69), 3PLS (38), 2QD9 (50), 2RKU (44), 3S95 (66), 3SOC (67), 3SXS (31), 4TPT (63), 3TXO (38), 4UAL (56), 1UWH (69), 2V62 (85), 4WB8 (56), 1WT6 (29), 3WZU (36), 4X2G (38), 4X7Q (60), 2XRW (56), 4YFI (154), 3ZBF (39), 3ZDU (40), 3ZH8 (118), 4ZZN (46) |
| **pazopanib** | 3ALO (37), 4ASD (43), 4BKJ (71), 2BUJ (83), 5C46 (92), 4C8B (80), 3CBL (41), 4CMT (45), 3COK (63), 3DTC (31), 4FGB (42), 1FMK (46), 4FVP (36), 2GK9 (58), 3HMI (29), 4HVS (40), 4ITJ (65), 2IVT (52), 2J7T (30), 4J8N (40), 2JAM (60), 4JS8 (35), 3LXL (37), 3LXP (29), 3MDY (100), 3MTL (42), 4O38 (75), 4OBO (58), 2OG8 (57), 2OGV (46), 2OO8 (37), 2P0C (63), 3Q6U (29), 2QLU (37), 2QNJ (80), 1RJB (41), 3S95 (65), 4TPT (63), 3UIU (88), 1UWJ (71), 3VN9 (54), 2W5A (33), 4WSQ (79), 1XBB (37), 2XRW (56), 3ZBF (39) | 3A7I (37), 3A99 (33), 4AAA (40), 2AC3 (32), 4AF3 (43), 4AGU (91), 4ASZ (40), 4AU8 (87), 4B9D (79), 4BF2 (52), 4BGQ (36), 4BKY (48), 3BLH (78), 1BO1 (106), 3CC6 (37), 2CMW (40), 3COM (85), 4CT2 (32), 3D7T (83), 4E5W (94), 3E7O (107), 1E8Z (22), 5EFQ (47), 3EQC (34), 2EU9 (44), 4EUU (81), 2EVA (44), 4EYJ (51), 4F6U (93), 4FKL (38), 4FR4 (253), 3FXZ (32), 3G2F (84), 4GV1 (51), 2H6D (31), 4HCU (30), 5HG8 (38), 4HYI (31), 4I4E (45), 4I5P (43), 2I6L (70), 4IC7 (123), 3IEC (88), 4IIR (116), 1J1B (105), 2JC6 (62), 4JPS (53), 4JSP (299), 4KIK (176), 3KN6 (65), 4KS7 (37), 3KVW (47), 4KWP (52), 4L00 (79), 3LM5 (31), 3M2W (37), 3MFS (33), 3MY0 (85), 4NW6 (36), 4OTH (40), 4OTP (33), 4OTW (44), 2OZO (82), 3P1A (37), 1P4O (84), 4PED (69), 4PF4 (38), 3PLS (38), 2QD9 (50), 2QOL (34), 4RFZ (40), 2RKU (44), 3SOC (67), 4TWC (67), 3TXO (38), 4U6R (63), 1UA2 (139), 4UAL (55), 2V62 (85), 3V8S (209), 2VWI (97), 1VZO (44), 2W4O (33), 2W96 (91), 1WAK (43), 4WB8 (56), 2WEL (55), 3WF7 (33), 2WNT (67), 4WNO (34), 2WQM (33), 1WT6 (29), 2WTK (237), 3WZU (34), 4X2G (38), 2X4F (79), 4X7Q (58), 1X8B (31), 1XJD (26), 2Y7J (119), 4Y73 (138), 2YCF (45), 4YFI (154), 4YHJ (121), 4YLK (43), 1Z57 (48), 3ZDU (38), 3ZH8 (117), 3ZON (34), 2ZV2 (38), 4ZZN (44) |

Supplementary Table 1: The pdb structures used to make the respective positive structural signatures (pockets extracted from the first chain) and negative signatures (pocket most similar to the bound pocket).
