## Supplemental Table 2 for "Identifying Novel Targets by using Drug-binding Site Signature: A Case Study of Kinase Inhibitors"

|  | **Sorafenib** | | **Sunitinib** | | **Dasatinib** | | **Imatinib** | | **Pazopanib** | | **Average** | |
| --- | --- | --- | --- | --- | --- | --- | --- | --- | --- | --- | --- | --- |
| **cut-off** | Sensitivity | Specificity | Sensitivity | Specificity | Sensitivity | Specificity | Sensitivity | Specificity | Sensitivity | Specificity | Sensitivity | Specificity |
| **0.85** | 0.81 | 0.33 | 0.63 | 0.67 | **0.02** | 0.99 | **0.03** | 0.94 | **0.04** | 0.95 | **0.31** | 0.78 |
| **1** | 0.94 | **0** | 0.94 | **0.12** | **0.21** | 0.76 | **0.35** | 0.73 | **0.24** | 0.75 | **0.54** | **0.33** |
| **1.15** | 1 | **0** | 1 | **0** | 0.75 | **0.23** | 0.65 | **0.27** | 0.65 | **0.24** | 0.81 | **0.15** |

Supplementary Table 2: Performance using positive signature alone: The positive structural signature alone is not sufficient in the case of kinase inhibitors to distinguish targets from non-targets as using different cut-offs the sensitivity or specificity becomes unacceptable. Values less than 60% are highlighted.
