## Supplemental Table 3 for "Identifying Novel Targets by using Drug-binding Site Signature: A Case Study of Kinase Inhibitors"

|  | **Positive Signature** | **Preservation ratio (%)** | **Negative Signature** | **Preservation ratio (%)** |
| --- | --- | --- | --- | --- |
| **sorafenib** | 31 | 50 | 90 | 60 |
| **sunitinib** | 93 | 60 | 60 | 60 |
| **dasatinib** | 52 | 60 | 111 | 65 |
| **imatinib** | 31 | 50 | 117 | 50 |
| **pazopanib** | 46 | 60 | 111 | 65 |

Supplementary Table 3: The number of structures used to make the positive and negative signatures and the redundancy cut-off for each signature.
