## Supplemental Table 4 for "Identifying Novel Targets by using Drug-binding Site Signature: A Case Study of Kinase Inhibitors"

| **PDB ID** | **Protein Name** | **Positive Score** | **Negative Score** | **Positive-Negative Score** | **Docking Score** | **Random Chance** |
| --- | --- | --- | --- | --- | --- | --- |
| 3GPH | Cytochrome P450 2E1 | 0.53 | 0.6 | -0.07 | -194.61 | <1% |
| 3MRK | MHC Class I (HLA A2) | - | - | - | -188.48 | <1% |
| 3KXC | Trafficking protein particle complex subunit 3 | 0.52 | 0.63 | -0.11 | -172.2 | 2% |
| 2BTS | Cell Division Protein Kinase 2 | 0.53 | 0.66 | -0.13 | -136.13 | 2% |
| 1T4V | Prothrombin | 0.51 | 0.71 | -0.20 | -104.62 | 2% |
| 3EBS | Cytochrome P450 2A6 | 0.53 | 0.89 | -0.36 | -100.57 | 4% |
| 1XTK | Probable ATP-dependent RNA helicase p47 | 0.50 | 0.78 | -0.28 | -74.49 | 6% |
| 1YB1 | 17-beta-hydroxysteroid dehydrogenase type XI | 0.53 | 0.59 | -0.06 | -74.06 | 6% |
| 3O2N | Ribosyldihydronicotinamide dehydrogenase [quinone] | 0.52 | 0.74 | -0.22 | -69.91 | 6% |
| 1ZVD | Smad ubiquitination regulatory factor 2 | 0.53 | 0.60 | -0.07 | -54.54 | 7% |
| 4IV2 | Estrogen receptor | 0.51 | 0.75 | -0.24 | -52.83 | 10% |

Supplementary Table 4: Top 10 predicted targets of Dasatinib, their PDB id, Score_positive_, Score_negative_, Score_positive_ – Score_negative_, docking score and the significance measure (random chance to obtain a better docking score).
