## Supplemental Table 5 for "Identifying Novel Targets by using Drug-binding Site Signature: A Case Study of Kinase Inhibitors"

| **PDB ID** | **Protein Name** | **Positive Score** | **Negative Score** | **Positive-Negative Score** | **Docking Score** | **Random Chance** |
| --- | --- | --- | --- | --- | --- | --- |
| 4HCV | Tyrosine-protein kinase ITK/TSK | 0.54 | 0.599 | -0.06 | -291.45 | <1% |
| 3MRK | MHC Class I (HLA A2) | - | - | - | -126.73 | 2% |
| 3CZY | Heme oxygenase 1 | 0.55 | 0.687 | -0.14 | -83.73 | 5% |
| 3ITK | UDP-glucose 6-dehydrogenase | 0.56 | 0.735 | -0.17 | -80.35 | 7% |
| 1PXO | Cell division protein kinase 2 | 0.58 | 0.614 | -0.03 | -79.15 | 5% |
| 4HHB | HEMOGLOBIN (DEOXY) (ALPHA CHAIN) | 0.57 | 0.624 | -0.05 | -68.99 | 9% |
| 2IO5 | ASF1A protein | 0.43 | 0.712 | -0.28 | -59.32 | 11% |
| 3Q93 | 7,8-dihydro-8-oxoguanine triphosphatase | 0.58 | 0.858 | -0.28 | -39.53 | 13% |
| 2I4J | Peroxisome proliferator-activated receptor gamma | 0.58 | 0.701 | -0.12 | -36.31 | 15% |
| 4AY9 | GLYCOPROTEIN HORMONES, ALPHA POLYPEPTIDE | 0.55 | 0.774 | -0.22 | -33 | 18% |

Supplementary Table 5: Top 10 predicted targets of Imatinib, their PDB id, Score_positive_, Score_negative_, Score_positive_ – Score_negative_, docking score and the significance measure (random chance to obtain a better docking score).
