## Supplemental Table 6 for "Identifying Novel Targets by using Drug-binding Site Signature: A Case Study of Kinase Inhibitors"

| **PDB ID** | **Protein Name** | **Positive Score** | **Negative Score** | **Positive-Negative Score** | **Docking Score** | **Random Chance** |
| --- | --- | --- | --- | --- | --- | --- |
| 1GZX | Hemoglobin Alpha Chain | 0.53 | 0.65 | -0.12 | -216.7 | 1% |
| 3BXN | MHC Class I (HLA B*1402) | - | - | - | -199.2 | 3% |
| 2FAI | Estrogen receptor | 0.52 | 0.74 | -0.22 | -154.7 | 7% |
| 1HAG | Prethrombin 2 | 0.56 | 0.80 | -0.24 | -99.6 | 9% |
| 2F6J | Bromodomain PHD finger transcription factor | 0.56 | 0.67 | -0.11 | -72.01 | 9% |
| 4DRA | Centromere protein S | 0.55 | 0.68 | -0.13 | -67.9 | 10% |
| 3FA6 | Cellular retinoic acid-binding protein 2 | 0.57 | 0.76 | -0.19 | -55.9 | 10% |
| 3IUY | Probable ATP-dependent RNA helicase DDX53 | 0.54 | 0.80 | -0.26 | -54.2 | 11% |
| 2P4Y | Peroxisome proliferator-activated receptor gamma | 0.53 | 0.74 | -0.21 | -43.7 | 13% |
| 4MCY | HLA class II histocompatibility antigen, DR alpha chain | 0.55 | 0.92 | -0.37 | -38.8 | 14% |

Supplementary Table 6: Top 10 predicted targets of Sunitinib, their PDB id, Score_positive_, Score_negative_, Score_positive_ – Score_negative_, docking score and the significance measure (random chance to obtain a better docking score).
