## Supplemental Table 7 for "Identifying Novel Targets by using Drug-binding Site Signature: A Case Study of Kinase Inhibitors"

| **PDB ID** | **Protein Name** | **Positive Score** | **Negative Score** | **Positive-Negative Score** | **Docking Score** | **Random Chance** |
| --- | --- | --- | --- | --- | --- | --- |
| 3MHJ | Tankyrase-2 | 0.56 | 0.942 | -0.382 | -154.43 | <1% |
| 1YHS | Proto-oncogene serine/threonine-protein kinase Pim-1 | 0.54 | 0.654 | -0.114 | -126.69 | <1% |
| 3QJC | Hemoglobin subunit alpha | 0.54 | 0.54 | -0.00 | -109.16 | <1% |
| 2HI4 | Cytochrome P450 1A2 | 0.52 | 0.677 | -0.157 | -105.97 | <1% |
| 2BEI | Diamine acetyltransferase 2 | 0.56 | 0.691 | -0.131 | -99.64 | <1% |
| 2ZK1 | Peroxisome proliferator-activated receptor gamma | 0.56 | 1.036 | -0.476 | -76.46 | 2% |
| 4K77 | Tyrosine-protein kinase JAK1 | 0.49 | 0.785 | -0.295 | -71.72 | 2% |
| 4DBS | Aldo-keto reductase family 1 member C3 | 0.55 | 0.746 | -0.196 | -58.03 | 2% |
| 4KC4 | Fucosylglycoprotein alpha-N-acetylgalactosaminyltransferase | 0.51 | 0.872 | -0.362 | -50.45 | 2% |
| 1MF8 | Calmodulin-Dependent Calcineurin A Subunit | 0.55 | 0.774 | -0.22 | -33 | 3% |
| 4MGC | Estrogen receptor | 0.57 | 0.76 | -0.19 | -43.23 | 3% |
| 4Y46 | Mitogen-activated protein kinase 10 | 0.57 | 0.81 | -0.24 | -24.05 | 7% |

Supplementary Table 7: Top 10 predicted targets of Pazopanib, their PDB id, Score_positive_, Score_negative_, Score_positive_ – Score_negative_, docking score and the significance measure (random chance to obtain a better docking score).
